# NOTCH3 p.R544C Drives Early Neurovascular Unit Dysfunction through Blood-Brain Barrier Failure, Complement Signalling and Metabolic Dysregulation

**DOI:** 10.64898/2026.11.17.767474

**Authors:** Jolene Wei Ling Lee, Zhi-Wei Zhang, Monika Chauhan, Xing Qi Teo, Alicia Qian Ler Tan, Tse Siew Lim, Hannah Wee, Zhen Kai Ngian, Laurie H Sanders, Christine Cheung, Zhong Pei, Adeline Su Lyn Ng, Qunxiang Ong, Eng-King Tan, Li Zeng

## Abstract

Cerebral small vessel disease is a major cause of stroke and vascular cognitive impairment, yet the mechanisms that initiate neurovascular dysfunction before irreversible brain injury remain poorly defined. CADASIL, caused by cysteine-altering NOTCH3 mutations, provides a genetically tractable model to resolve these early events. Here, we investigated the Asian-enriched NOTCH3 p.R544C variant and tested whether early disease emerges through interconnected blood-brain barrier (BBB), inflammatory, and metabolic dysfunction.

NOTCH3 p.R544C iPSC-derived endothelial cells showed impaired angiogenesis and migration, reduced barrier resistance, disrupted tight-junction organisation, and altered vascular, adhesion, and extracellular-matrix programmes. In multicellular BBB-like microtissues, these defects propagated across the neurovascular unit, producing disorganised vascular networks, enlarged luminal structures, and increased permeability, establishing BBB failure as an early disease layer.

Single-cell transcriptomics revealed coordinated loss of endothelial BBB transport and efflux programmes with astrocytic complement, innate immune, and matrix-remodelling activation. Secretome profiling and patient-plasma measurements converged on complement- and matrix-associated mediators, including C1S and CLU, identifying complement-matrix activation as a second interconnected layer. Importantly, combined cilostazol and isosorbide mononitrate treatment improved barrier function and reduced selected complement- and matrix-associated proteins in mutant microtissues, supporting pharmacological reversibility of components of this early disease programme.

Seahorse analysis further demonstrated reduced mitochondrial respiratory capacity associated with NOTCH3 p.R544C. *In vivo*, three-month-old mutant mice showed altered cerebral pyruvate metabolism and early microvascular abnormalities, whereas six-month-old mice developed more prominent extracellular-matrix, inflammatory, and gliovascular changes, supporting an age-dependent transition from early metabolic dysregulation to broader neurovascular remodelling.

Together, these findings define NOTCH3 p.R544C CADASIL as an early multicellular neurovascular disorder organised across three interconnected mechanistic layers: BBB failure, complement-matrix signalling, and metabolic dysregulation, while identifying early BBB-centred dysfunction as a potentially tractable state before advanced cerebrovascular injury develops.

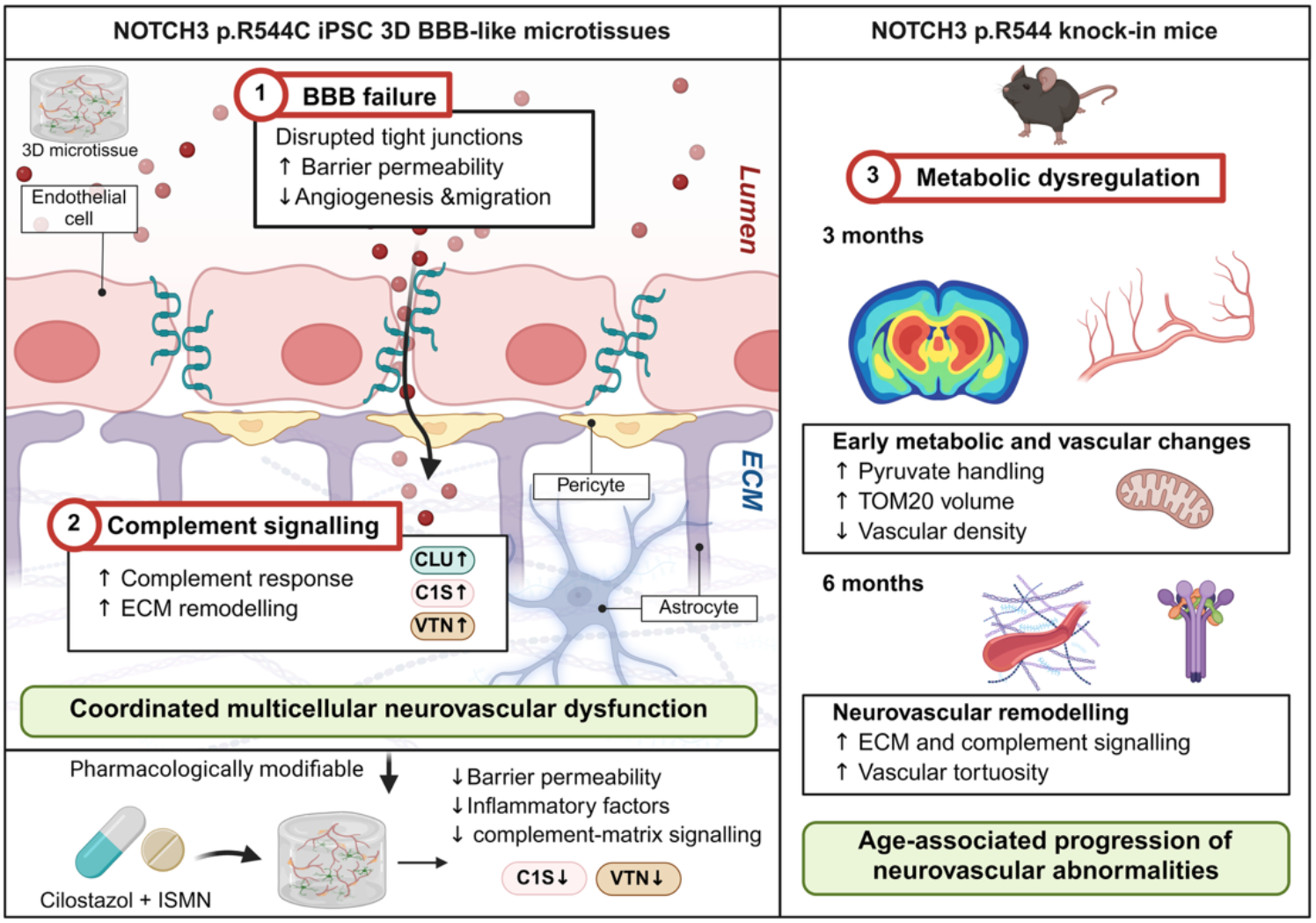

**HIGHLIGHTS:**

- NOTCH3 p.R544C establishes an intrinsic endothelial disease state that initiates BBB failure, impairing angiogenesis, tight-junction organisation, and barrier function.
- BBB failure propagates across the neurovascular unit, producing abnormal multicellular organisation, vascular remodelling, and increased permeability.
- Single-cell profiling identifies complement signalling as a second mechanistic layer, linking endothelial BBB programme loss with astrocytic complement-matrix and innate immune responses.
- Cilostazol and isosorbide mononitrate improve BBB function and attenuate complement-matrix abnormalities, supporting the reversibility of this early neurovascular disease programme.
- Metabolic dysregulation forms a third mechanistic layer, with impaired mitochondrial respiration and early metabolic abnormalities preceding progressive neurovascular and inflammatory remodelling *in vivo*.

## Introduction

### Early neurovascular changes in cerebral small vessel disease

Cerebral small vessel disease (CSVD) encompasses a heterogeneous group of disorders affecting the brain microvasculature and is a major cause of stroke, cognitive impairment and dementia^1,2^. Although radiological manifestations such as white matter hyperintensities, lacunes and cerebral microbleeds are well characterised, these features represent relatively late consequences of a disease process that likely begins years earlier within the cerebral microcirculation^3–5^. Increasing evidence places dysfunction of the neurovascular unit at the centre of this early phase. Cerebral endothelial cells maintain blood-brain barrier (BBB) integrity, regulate nutrient transport and vascular tone, and coordinate signalling with pericytes, astrocytes and other neural cells^6^. Disruption of these functions can alter vascular permeability and tissue homeostasis before overt structural injury becomes apparent and have been implicated in other neurodegenerative diseases^7–19^. Defining these early and potentially reversible events remains challenging in sporadic CSVD, in which ageing, vascular risk factors and genetic susceptibility frequently coexist.

### CADASIL as a genetically defined model of small vessel disease

Cerebral autosomal dominant arteriopathy with subcortical infarcts and leukoencephalopathy (CADASIL) provides a genetically defined system in which to dissect the initiating mechanisms of CSVD^20–25^. The disorder is caused by pathogenic variants in NOTCH3, most of which alter the number of cysteine residues within epidermal growth factor-like repeats of the receptor extracellular domain^26^. NOTCH3 is highly expressed in vascular smooth muscle cells and pericytes and is required for arterial development and vascular homeostasis^27^. A pathological hallmark of CADASIL is progressive accumulation of NOTCH3 extracellular domain within vessel walls, together with granular osmiophilic material and extracellular-matrix proteins^26,28–30^. These changes are accompanied by mural-cell degeneration, vascular remodelling and progressive white matter injury, establishing mural-cell dysfunction as a central feature of disease pathogenesis.

### Endothelial and BBB dysfunction in CADASIL

However, mural-cell degeneration alone does not fully explain how mutant NOTCH3 initiates brain injury. NOTCH3 extracellular-domain deposition can be detected before advanced clinical disease and extends beyond larger arteries and arterioles into the cerebral microvasculature^29,31^. Experimental and human studies increasingly implicate endothelial dysfunction, impaired neurovascular signalling and BBB disruption in CADASIL and other forms of CSVD^12,32,33^. Human induced pluripotent stem cell and cell line models carrying CADASIL-associated NOTCH3 variants have demonstrated impaired endothelial barrier properties and abnormal endothelial-mural-cell interactions^34–38^, raising the possibility that BBB dysfunction represents an early component of disease rather than a secondary consequence of established vascular degeneration. The unresolved question is therefore whether mutant NOTCH3 establishes an early endothelial disease state that subsequently propagates across the neurovascular unit before irreversible structural injury develops.

### Endothelial-astrocyte signalling, complement activation and neurovascular propagation

This question is particularly important because endothelial dysfunction is unlikely to act in isolation. Astrocytic endfeet closely ensheath the cerebral microvasculature and regulate endothelial junctional integrity, basement membrane organisation, water and ion homeostasis, neurovascular coupling and inflammatory signalling^39,40^. We therefore reasoned that an initial endothelial barrier defect could perturb reciprocal signalling within the neurovascular unit and induce secondary astrocytic responses that amplify vascular dysfunction. Complement signalling is a plausible mediator of this process because locally produced complement components can regulate innate immune activation, extracellular-matrix remodelling and vascular integrity^41–43^. Whether endothelial dysfunction in CADASIL is coupled to astrocytic complement activation, and whether this interaction contributes to early neurovascular pathology, remain unknown. Resolving these responses at single-cell resolution could clarify how mutant NOTCH3 reshapes signalling across distinct neurovascular cell populations.

### Neurovascular dysfunction and cerebral metabolic homeostasis

Beyond maintaining barrier integrity, the neurovascular unit plays an essential role in cerebral metabolic homeostasis by coordinating substrate delivery, endothelial transport, vascular perfusion and metabolic interactions between vascular and neural cells^44^. Disturbances in these processes may alter cerebral energy metabolism before overt structural injury becomes apparent. Mitochondrial dysfunction and impaired metabolic regulation have increasingly been implicated in CSVD, where altered energy availability and oxidative stress may contribute to vascular dysfunction and tissue vulnerability^8,45,46^. Previous studies have also reported mitochondrial abnormalities in experimental models of CADASIL, suggesting that mutant NOTCH3 may affect cellular bioenergetics beyond its established effects on vascular structure ^45–48^. However, whether metabolic alterations occur during the early stages of NOTCH3-associated disease, and how they relate to BBB dysfunction and subsequent neurovascular remodelling, remains poorly understood. Defining these early metabolic changes alongside vascular and inflammatory responses may provide a more integrated understanding of disease initiation and progression.

### NOTCH3 p.R544C in East Asian populations

These questions are especially relevant to the NOTCH3 p.R544C variant. In contrast to the predominance of NOTCH3 variants within epidermal growth factor-like repeats 1-6 in many European CADASIL cohorts^20,49^, p.R544C is one of the most common variant globally^25^ and highly enriched in East Asian populations^24,25,50–57^. It accounts for a large proportion of CADASIL pedigrees reported in Taiwan and has also been identified in population-based Taiwanese and Singapore Chinese cohorts^51,52^. Carriers show increased risk of small-vessel occlusion stroke, yet clinical penetrance and disease severity vary considerably^21,50,51,56,58–64^. This combination of relatively high population frequency and heterogeneous clinical expression makes p.R544C a particularly informative model in which to investigate early neurovascular mechanisms that precede overt cerebrovascular injury.

### Unresolved mechanisms in multicellular human neurovascular models

Despite its regional importance, the early cellular consequences of NOTCH3 p.R544C remain incompletely defined. Recent work supports endothelial dysfunction associated with this variant^47,65^, but it remains unclear whether these abnormalities persist in a multicellular human neurovascular environment, which neighbouring cell populations contribute to the resulting phenotype, and whether these cellular changes converge with molecular alterations detectable in patients and *in vivo* models. Conventional two-dimensional cultures cannot reproduce lumen formation, three-dimensional (3D) vascular architecture or reciprocal signalling among endothelial cells, pericytes and astrocytes. Conversely, animal models provide physiological context but make it difficult to distinguish primary cellular responses from secondary changes that accumulate during disease progression.

### Study hypothesis and cross-scale mechanistic strategy

Here, we tested the hypothesis that NOTCH3 p.R544C establishes an early endothelial barrier defect that propagates across the neurovascular unit through altered endothelial-glial signalling, including complement- and extracellular-matrix-associated pathways, before advanced structural vascular pathology develops. We combined an isogenic human induced pluripotent stem cell system with 3D BBB-like microtissues, single-cell transcriptomics, patient plasma profiling and a NOTCH3 p.R544C mouse model. This cross-model approach allowed us to examine the sequence from intrinsic endothelial dysfunction to multicellular neurovascular dysregulation and its *in vivo* correlates. Our findings identify a disease state characterised by impaired endothelial barrier function and loss of BBB-associated endothelial programmes together with astrocytic complement activation, complement- and extracellular-matrix-associated changes in patient plasma, and early metabolic and gliovascular abnormalities *in vivo*. Pharmacological improvement of barrier function in mutant microtissues further indicates that components of this early neurovascular phenotype remain modifiable. Collectively, these results support a model in which NOTCH3 p.R544C induces an early, multicellular neurovascular disorder centred on BBB dysfunction and altered endothelial-glial signalling, revealing a potentially tractable disease phase before irreversible cerebrovascular injury develops.

## Results

### NOTCH3 p.R544C establishes an intrinsic endothelial disease state characterised by impaired angiogenesis and barrier dysfunction

To determine whether NOTCH3 p.R544C (MT) directly alters endothelial function, we compared patient-derived iPSC endothelial cells carrying the variant with an isogenic corrected control. Both lines reproducibly acquired endothelial morphology and expressed vascular markers, enabling mutation-associated phenotypes to be assessed in a matched genetic background.

We first examined angiogenic network formation. Corrected ECs formed interconnected networks on basement membrane matrix, whereas MT ECs generated visibly less extensive and less connected structures (**Fig. 1A**). Quantitative image analysis showed reductions in total vessel length and number of junctions and vessel area (**Fig. 1B,F,G**), together with increased vessel diameter (**Fig. 1C**). These measures indicate impaired network formation rather than a change confined to one angiogenic parameter.

**Figure 1.**
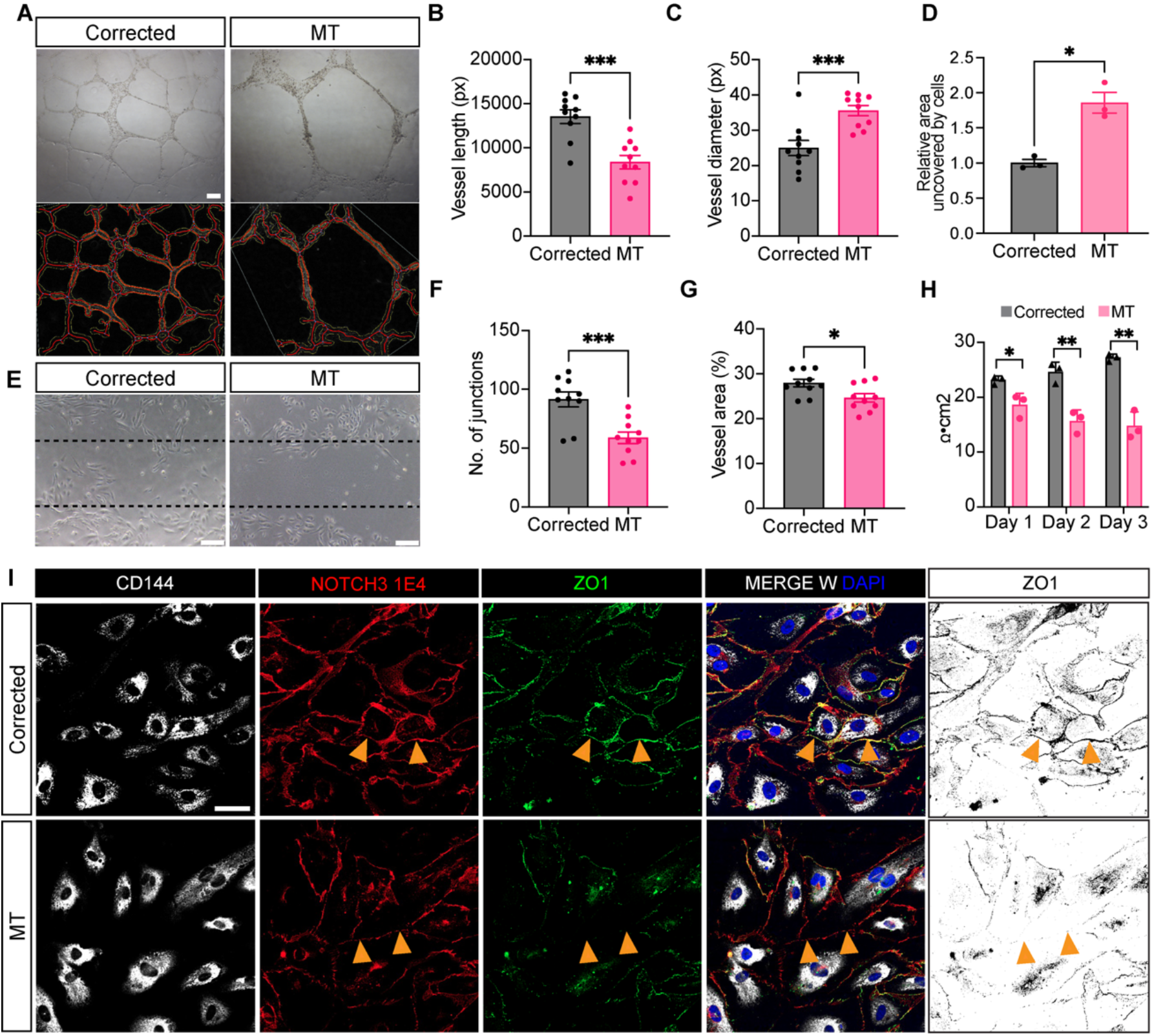
NOTCH3 p.R544C establishes an intrinsic endothelial disease state characterised by impaired angiogenesis and barrier dysfunction. **(A)** Representative angiogenic network images of isogenic corrected and MT iPSC-derived endothelial cells. n = 10 per condition, scale bar = 200μm. **(B,C,F,G)** Quantification of vessel length **(B)**, vessel diameter **(C)**, number of junctions **(F)** and vessel area **(G)**. **(D)** Quantification of relative uncovered area following scratch formation, with corresponding representative images in **(E)**. n = 3 per condition, scale bar = 100μm. **(H)** Transendothelial electrical resistance over three days in corrected and MT endothelial cell monolayers. **(I)** Representative CD144 (white), NOTCH3 extracellular domain (1E4) (red) and ZO1 (green) immunostaining, including merged images with DAPI (blue) and processed ZO1 binary images; arrowheads indicate selected junctional regions. Scale bar = 50μm. Data are presented as mean ± SEM, *p < 0.05, ***p < 0.001 by Welch’s t-test.

Endothelial migration was assessed independently using a scratch assay. At 18 h, MT monolayers retained a larger uncovered area, approximately 87% higher, than corrected controls (**Fig. 1D,E**). This finding is consistent with reduced wound closure and complements the network-formation phenotype.

We next asked whether p.R544C-associated differences extended to the formation of an endothelial barrier. Transendothelial electrical resistance remained significantly lower in MT monolayers during the three-day measurement period, with an approximately 40% reduction on day 3 compared with corrected endothelial cells (15.47 versus 25.91 Ω·cm²; **Fig. 1H**). Immunostaining showed irregular, less continuous localisation of the tight junction-associated protein ZO1 at MT cell-cell borders, in contrast to the more continuous junctional pattern in corrected ECs (**Fig. 1I**).

As astrocytes provide an important source of neurovascular support, we further examined whether the variant was associated with overt changes in astrocyte morphology under monolayer conditions. GFAP staining and Sholl analysis showed broadly similar process architecture in corrected and MT astrocytes, with overlapping radial intersection profiles and distributions of intersection number, branch number, branch length and ramification index (**Supplementary Fig. 2A-F**).

Together, these findings define a cell-intrinsic endothelial disease state characterised by impaired angiogenesis, migration, and barrier integrity. This prompted us to test whether the endothelial defect propagates across the neurovascular unit when MT cells are incorporated into a multicellular BBB environment.

### NOTCH3 p.R544C disrupts multicellular BBB organisation and induces maladaptive vascular remodelling

Endothelial barrier properties *in vivo* are shaped by reciprocal interactions with astrocytes, pericytes and the surrounding extracellular matrix. We therefore generated 3D BBB-like microtissues by combining iPSC-derived endothelial cells, astrocytes and pericytes (**Supplementary Fig. 1**) within a fibrinogen/thrombin mix (**Fig. 2A**). By day 7, both corrected and MT microtissues formed interconnected vascular networks, allowing structural and functional consequences of the mutation to be assessed in a common multicellular environment.

**Figure 2.**
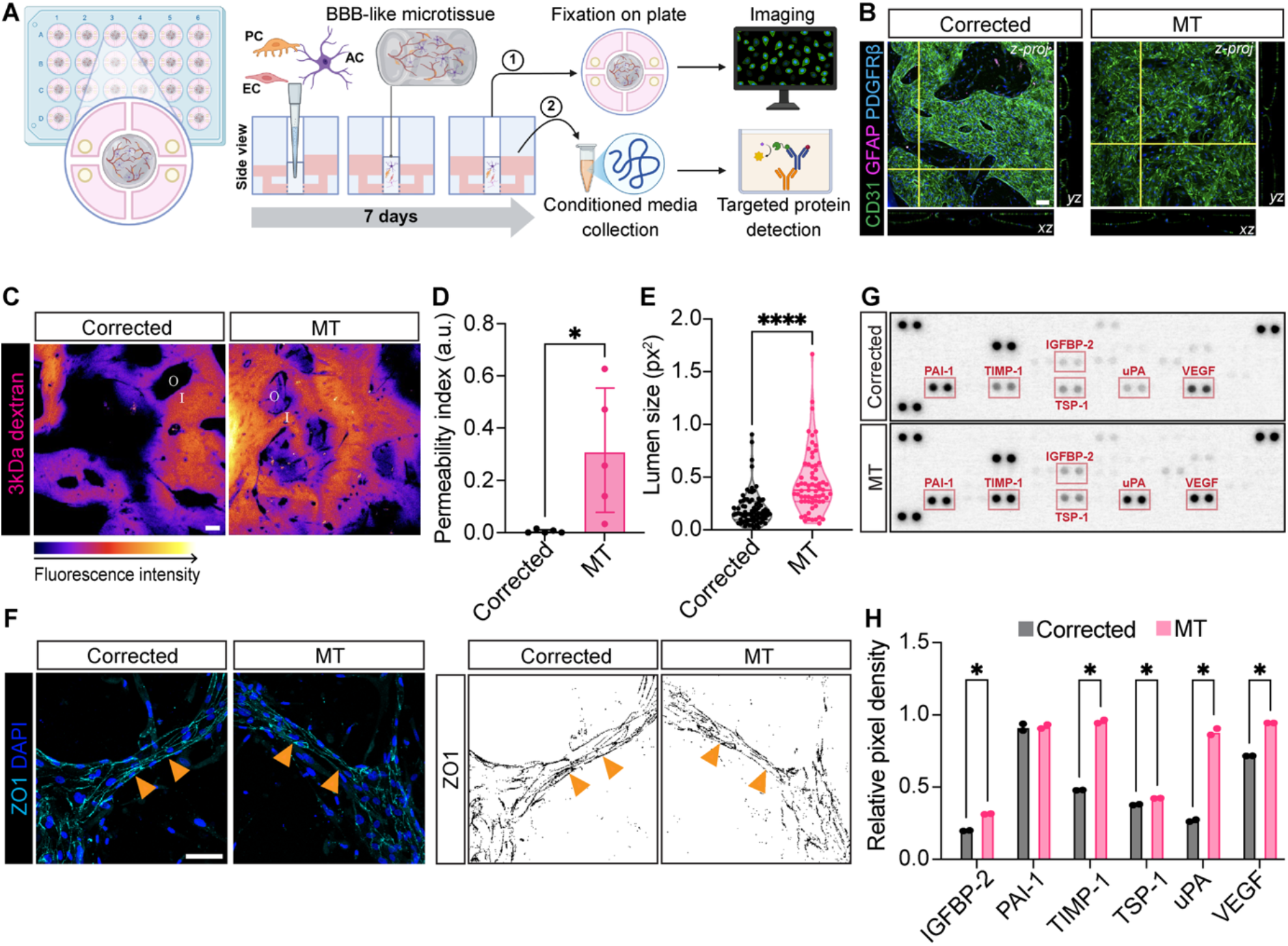
NOTCH3 p.R544C disrupts multicellular BBB organisation and drives maladaptive neurovascular remodelling. **(A)** Overview of microtissue assembly, imaging and conditioned-medium collection. **(B)** Representative CD31 (green), GFAP (magenta) and PDGFRβ (cyan) staining of corrected and MT microtissues, including orthogonal views. **(C)** Representative fluorescence-intensity maps following 3 kDa dextran perfusion. **(D)** Permeability index, calculated as extravascular divided by intravascular fluorescence intensity. **(E)** Quantification of CD31-defined lumen size. **(F)** Representative ZO1 (cyan) staining merged with DAPI (blue) and binary images of ZO1 vascular junctions. **(G)** Representative angiogenesis proteome arrays and **(H)** relative spot intensities for selected proteins in conditioned medium. Five microtissues per condition contributed to permeability and lumen analysis. Individual lumens are nested within microtissues. Secretome samples for the angiogenesis array were pooled from three microtissues per condition. Data are presented as mean ± SEM, *p < 0.05, ***p < 0.001 by Welch’s t-test. Scale bar = 50μm.

Although MT vessels remained perfusable, their organisation differed markedly from corrected microtissues (**Fig. 2B,C**). Following perfusion with 3 kDa fluorescent dextran, corrected vessels largely retained the tracer within the vascular compartment, whereas MT networks showed extensive extravascular fluorescence. The permeability index ratio increased an approximately 68-fold increase in relative tracer leakage (**Fig. 2D**). As both conditions formed patent vessels, the permeability phenotype reflects loss of barrier containment rather than failure of lumen formation.

Vascular architecture was altered in parallel. CD31-based analysis demonstrated enlarged luminal regions in MT microtissues, with the lumen-size measurement increasing approximately 2.4-fold (**Fig. 2E**). Junctional organisation was also compromised: ZO1 staining was more continuous along corrected microvessels, whereas MT networks showed irregular and fragmented localisation (**Fig. 2F**). These findings recapitulated the endothelial monolayer phenotype but within a setting in which endothelial cells were exposed to astrocytic, pericytic and matrix-derived signals.

We next asked whether structural barrier failure was accompanied by changes in the extracellular signalling environment. Angiogenesis-focused profiling of conditioned media identified increased abundance of several factors associated with vascular stress, angiogenesis and matrix turnover in MT microtissues (**Fig. 2G,H**). These included VEGF, uPA, TIMP-1, thrombospondin-1, uPA and IGFBP-2. The coexistence of increased vascular-remodelling factors with poorly organised and hyperpermeable networks argues against productive angiogenesis and instead suggests a maladaptive response to vascular stress. We therefore used single-cell transcriptomics to resolve the cellular origin of these changes within the MT neurovascular unit.

### Single-cell profiling reveals coordinated endothelial BBB programme loss and astrocytic complement-matrix activation with relevance to human CADASIL

To resolve transcriptional responses within the multicellular model, we performed single-cell RNA sequencing of corrected and MT microtissues. The integrated dataset contained four annotated cell populations corresponding to endothelial cells, astrocytes, pericytes and glial progenitor-like cells (**Fig. 3A-C**). Additional sample-level embeddings, quality-control measures and cluster-specific analyses are provided in **Supplementary Fig. 3**. This approach enabled comparison of mutation-associated responses across the constituent neurovascular populations.

**Figure 3.**
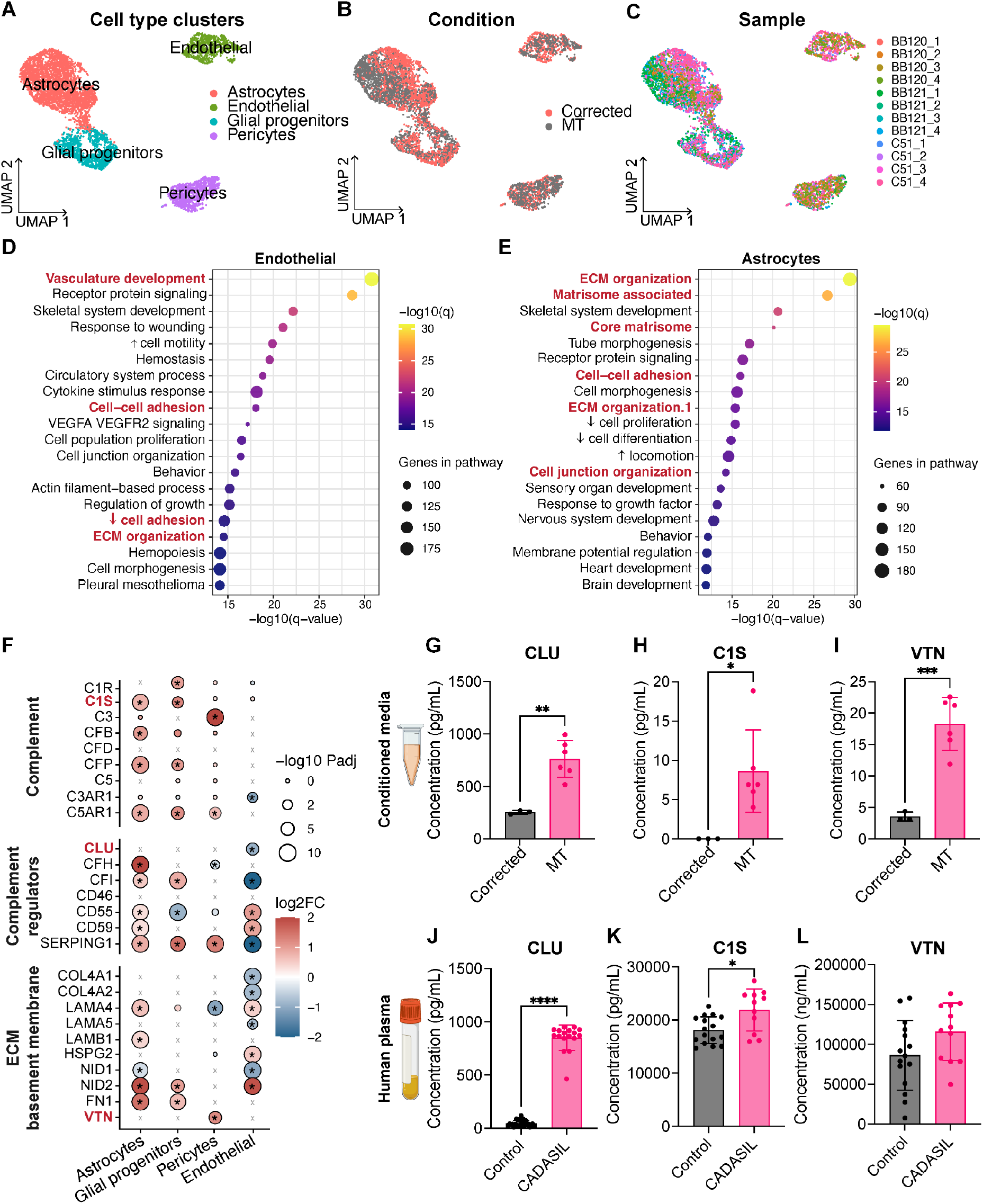
Single-cell profiling reveals coordinated loss of endothelial BBB programmes and activation of astrocytic complement signalling. **(A-C)** UMAP embeddings coloured by annotated cell type **(A)**, experimental condition **(B)** and individual sample **(C)**. 4 corrected and 8 MT microtissues were used for single-cell RNA sequencing. **(D,E)** Selected pathway-enrichment analyses in endothelial cells **(D)** and astrocytes **(E)**. Plot symbols encode pathway-gene number and enrichment significance as indicated. **(F)** Cell-type-specific expression differences in selected complement components and regulators and extracellular matrix-associated genes; colour indicates log2 fold change and size indicates adjusted significance. **(G-I)** CLU **(G)**, C1S **(H)** and VTN **(I)** concentrations in corrected and MT microtissue-conditioned medium. **(J-L)** CLU (29 control and 17 CADASIL samples) **(J)**, C1S (16 control and 11 CADASIL samples) **(K)** and VTN (15 control and 12 CADASIL samples) **(L)** concentrations in control and CADASIL plasma. Data are presented as mean ± SEM, *p < 0.05, **p < 0.01, ***p < 0.001, ****p < 0.0001 by Welch’s t-test.

Rather than producing a uniform transcriptional response, NOTCH3 p.R544C induced distinct but coordinated changes across neurovascular cell types. Endothelial pathway analysis highlighted vascular development, receptor signalling and processes related to cellular organisation (**Fig. 3D**). These pathways complemented the experimentally observed abnormalities in endothelial network formation and barrier properties. Astrocytes showed prominent enrichment of extracellular matrix organisation, matrisome and adhesion-associated terms (**Fig. 3E**), suggesting that the MT phenotype is also accompanied by changes in glial and extracellular programmes.

Glial progenitor cells exhibited alterations in developmental and morphogenetic pathways, including tube morphogenesis, cell junction organisation, extracellular matrix organisation, neuronal projection development and processes regulating cell proliferation and differentiation. In contrast, pericytes showed prominent enrichment of vascular and matrix-associated pathways, including vasculature development, extracellular matrix organisation, cell-cell adhesion and chemotaxis, alongside growth factor, MAPK and receptor tyrosine kinase signalling (**Supplementary Fig. 4**).

Focused analysis of curated functional gene modules further highlighted distinct transcriptional responses across cell types (**Supplementary Fig. 5**). Endothelial cells showed reduced expression of genes associated with BBB transport and efflux, tight junction organisation and complement-associated signalling, whereas astrocytes exhibited increased expression of complement, extracellular matrix and inflammatory response-associated genes. Glial progenitor-like cells and pericytes displayed additional changes in vascular signalling, extracellular matrix and complement-associated programmes, reinforcing the heterogeneous transcriptional response within MT microtissues. These findings suggest broader transcriptional changes involving cellular organisation, vascular support and extracellular matrix regulation across additional neurovascular populations.

Studies have implicated the role of the complement system in neuroinflammation and BBB dysfunction^41,42,66,67^. As such, we next examined complement-associated genes in the different cell populations. Complement-associated transcriptional changes varied across neurovascular cell populations. C1R and C1S were upregulated in astrocytes and glial progenitor-like cells, whereas C3 showed a prominent increase in pericytes. Among complement regulators, CFH was predominantly upregulated in astrocytes, while SERPING1 showed increased expression in astrocytes, glial progenitor-like cells and pericytes, but was downregulated in endothelial cells (**Fig. 3F**). These findings highlight distinct cell-type-associated complement transcriptional responses within MT microtissues, with astrocytes showing particularly broad changes in complement components and regulatory genes. (**Fig. 3F**). The same analysis highlighted alterations in extracellular matrix and basement membrane-associated genes across neurovascular populations. These results support altered complement-related transcription and matrix-associated signalling in the MT microtissue environment.

We selected CLU, C1S and VTN for protein-level follow-up to determine whether selected complement- and matrix-associated signals could also be detected outside the single-cell dataset. CLU and C1S are associated with complement regulation and signalling, whereas VTN has an established extracellular matrix role and has previously been reported in CADASIL-affected vessel walls and *in vitro* studies^30,68–70^. This histopathological association provided a particular rationale for including VTN even though changes in circulating concentrations may not mirror local vascular deposition. ELISA measurements showed higher concentrations of CLU, C1S and VTN in conditioned medium from MT microtissues than from corrected cultures (**Fig. 3G-I**).

To investigate relevance to human disease, we measured the same proteins in plasma from individuals with CADASIL and controls. CLU and C1S concentrations were higher in the CADASIL group (**Fig. 3J,K**). VTN showed a difference in group means but did not reach statistical significance (**Fig. 3L**). This further illustrates that the BBB-like microtissues recapitulate disease-associated factors.

Together, single-cell transcriptional profiling, microtissue secretion, and patient-plasma measurements converge on a complement-matrix programme associated with early neurovascular dysfunction. These findings extend the primary endothelial BBB defect into a broader multicellular disease state and identify astrocytic complement-matrix activation as a second mechanistic layer of NOTCH3 p.R544C pathology.

### Cilostazol and isosorbide mononitrate restore BBB function and attenuate complement-matrix-associated signals

The formation of vascular structures despite substantially increased permeability raised the possibility that at least part of the MT microtissue phenotype could be modified pharmacologically. We therefore examined combined cilostazol and isosorbide mononitrate (ISMN), two agents with complementary vascular actions^71–74^ being evaluated for combination treatment in CSVD^75–77^. Corrected and MT microtissues received vehicle or treatment as described in Methods. After 3 kDa dextran perfusion, treated MT networks showed less diffuse extravascular fluorescence than vehicle-treated MT networks (**Fig. 4A**). Quantification confirmed a reduction in the permeability index under treatment (**Fig. 4B**), indicating improved barrier-associated tracer containment in this experimental system.

**Figure 4.**
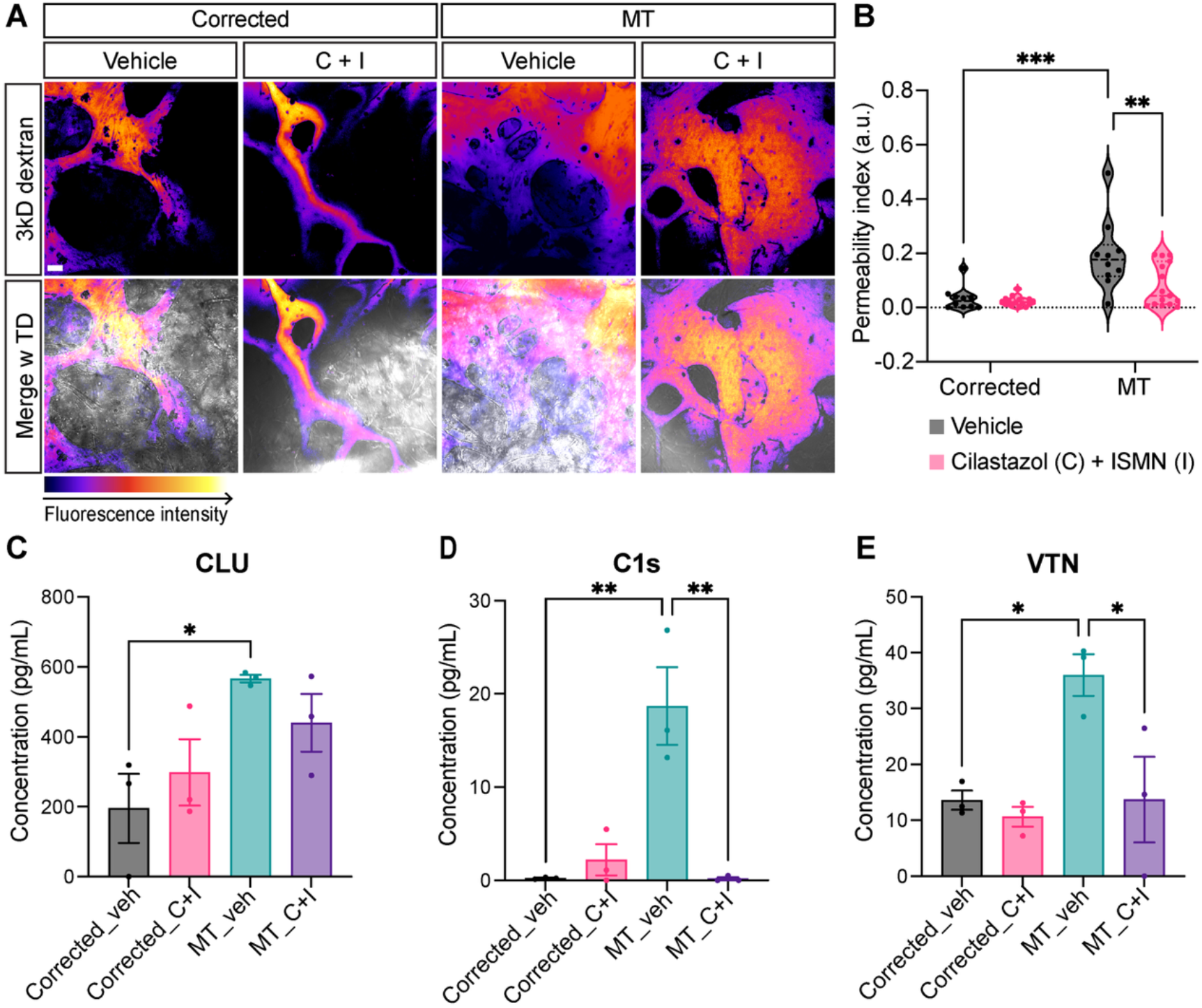
Cilostazol and isosorbide mononitrate restore BBB function and suppress the complement-matrix disease programme. **(A)** Representative fluorescence maps and transmitted-light overlays following 3 kDa dextran perfusion in corrected and MT BBB-like microtissues treated with vehicle or 1 μM cilostazol + 10 μM ISMN (C + I) for 7 days. **(B)** Permeability index by treatment group. **(C)** CLU, **(D)** C1S and **(E)** VTN concentrations in conditioned medium. Scale bar = 50μm. Data are presented as mean ± SEM, *p < 0.05, **p < 0.01, ***p < 0.001 by ordinary one-way ANOVA.

We then investigated whether the functional change coincided with differences in selected complement- and matrix-associated proteins. CLU varied among conditions but was not consistently reduced by treatment in MT microtissues (**Fig. 4C**). By contrast, C1S and VTN concentrations were lower in treated than in vehicle-treated MT cultures (**Fig. 4D,E**). Broader secretome profiling further showed treatment-dependent modulation of inflammatory, extracellular-matrix, and angiogenic factors, including IL-8, GDF15, CXCL1, ANGPT2, MMP9, osteopontin, VEGF, IGFBP-2, and IGFBP-3 (**Supplementary Fig. 6**). These changes support broader remodelling of the MT secretory environment while reinforcing the selective nature of the treatment response.

Together, the reduction in vascular permeability and attenuation of C1S and VTN demonstrate that both the functional barrier defect and selected components of the complement-matrix-associated disease state are pharmacologically modifiable. This reversibility supports the existence of a potentially tractable therapeutic window before more advanced structural pathology develops.

### NOTCH3 p.R544C impairs mitochondrial respiratory capacity and cellular bioenergetics

The human BBB studies identified endothelial dysfunction together with transcriptional changes involving metabolism and cellular stress. We therefore examined whether MT alters mitochondrial function in an independent cell system. HeLa-mKeima and HEK293T cells expressing wild-type (WT) or MT NOTCH3 were analysed using mitochondrial imaging and extracellular flux measurements respectively (**Fig. 5A**).

**Figure 5.**
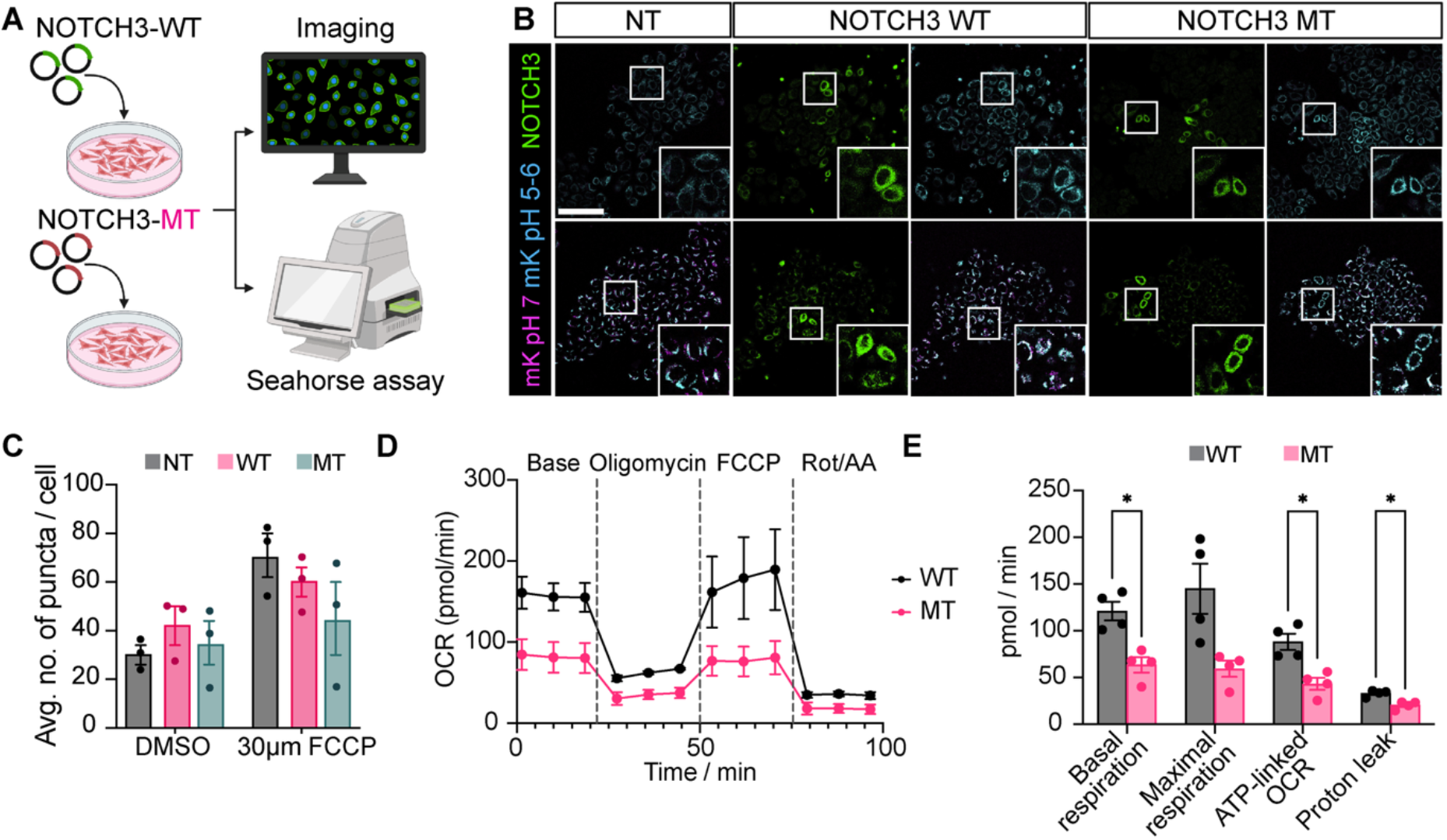
NOTCH3 p.R544C impairs mitochondrial respiratory capacity and cellular bioenergetics. **(A)** Study outline for WT and MT NOTCH3 expression with mitochondrial imaging and Seahorse analysis. **(B)** Representative mitochondrial reporter (pH 7 in magenta; pH 5-6 in cyan and NOTCH3 in green) images of non-transfected (NT), WT and MT NOTCH3-expressing HeLa-mKeima cells after vehicle or FCCP exposure. **(C)** Number of reporter-positive puncta per cell. n = 3 per condition. Scale bar = 50μm. **(D)** Oxygen consumption rate (OCR) during the mitochondrial stress test in HEK293T cells expressing WT or MT NOTCH3, following oligomycin, FCCP and rotenone/antimycin A exposure. **(E)** Quantification of basal respiration, maximal respiration, ATP-linked respiration and proton leak. n = 4 per condition. Data are presented as mean ± SEM *p < 0.05 by Welch’s t-test.

A pH-sensitive mitochondrial reporter in HeLa cells was used to examine mitochondrial puncta under basal conditions and following carbonyl cyanide *p*-trifluoromethoxyphenylhydrazone (FCCP) exposure (**Fig. 5B,C**). Although FCCP altered the overall punctate signal, the difference between WT and MT NOTCH3-expressing cells was modest and did not provide strong evidence for a genotype-dependent change in this readout.

In contrast, mitochondrial respiration was consistently reduced in cells expressing MT NOTCH3. Seahorse analysis showed lower oxygen consumption across the mitochondrial stress test (**Fig. 5D**). Quantification demonstrated reduced basal respiration and ATP-linked oxygen consumption in MT cells, together with reduced proton leak (**Fig. 5E**). Maximal respiratory capacity was also lower in MT cultures, although this difference was less pronounced.

Together, these data provide functional evidence that NOTCH3 p.R544C alters mitochondrial energetic capacity. This mitochondrial phenotype complements the BBB and complement-matrix abnormalities identified in the human models and prompted us to determine whether metabolic dysregulation emerges early during disease progression *in vivo*.

### NOTCH3 p.R544C drives an age-dependent shift from metabolic dysregulation to neurovascular and inflammatory remodelling *in vivo*

To determine the cellular findings were reflected *in vivo*, we analysed mice carrying the ROSA26-targeted human NOTCH3 p.R544C allele (NOTCH3^KI^) and controls (WT) at three and six months of age (**Fig. 6A**). These groups represent separate age-stratified cohorts; comparisons between ages therefore describe age-associated differences rather than within-animal longitudinal progression.

**Figure 6.**
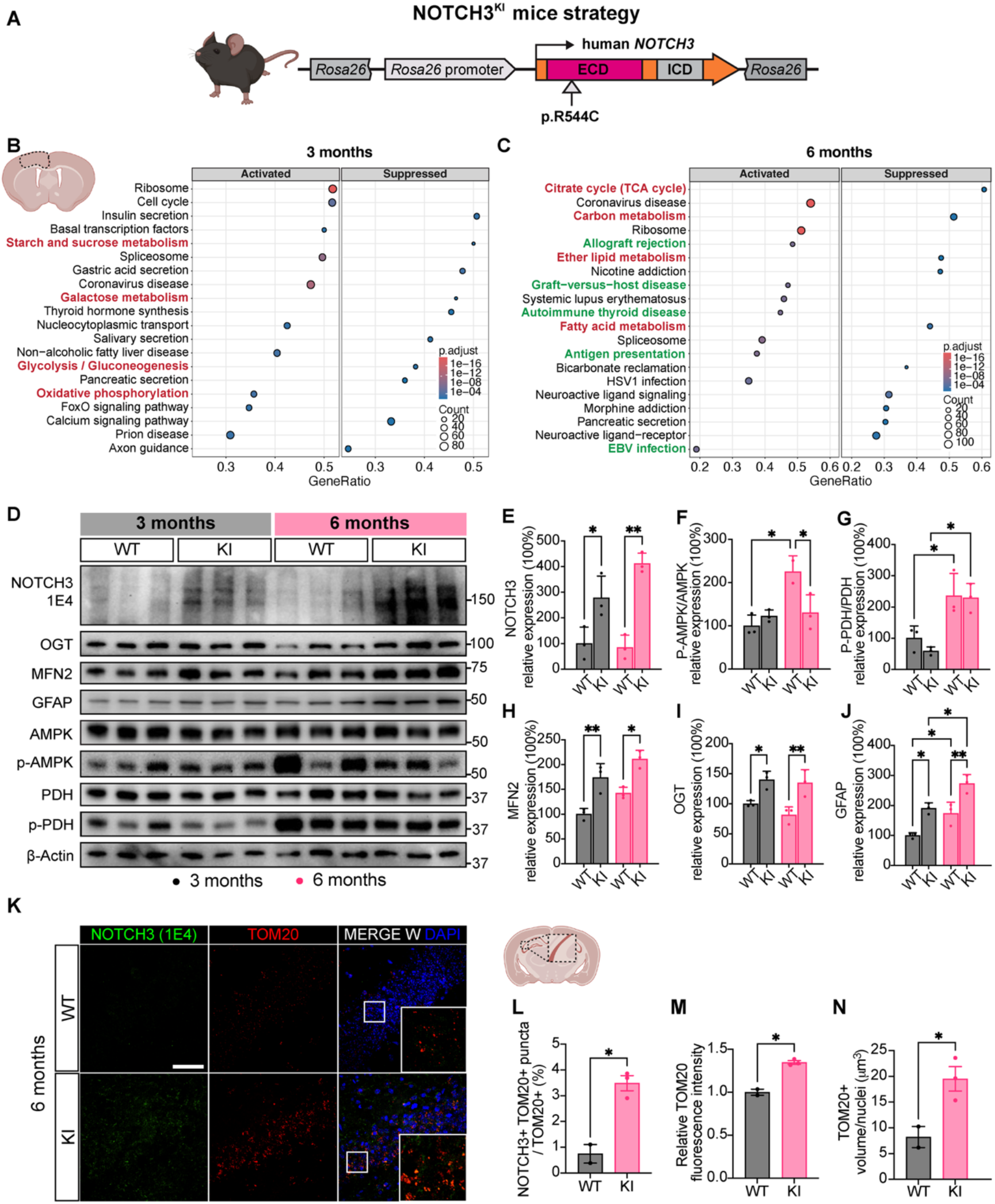
NOTCH3 p.R544C drives an age-dependent transition from early metabolic dysfunction to neurovascular and inflammatory remodelling *in vivo*. **(A)** Schematic of the ROSA26-targeted human NOTCH3 p.R544C allele for the generation of NOTCH3 p.R544C mice (NOTCH^KI^). Experimental comparisons are between NOTCH3^KI^ (KI in figure) and WT mice. **(B,C)** Ranked KEGG gene-set enrichment of bulk RNA sequencing of mouse cortical tissue at three **(B)** and six **(C)** months, displayed as positively and negatively enriched pathways. n = 2 mice per group. **(D)** Representative brain immunoblots for NOTCH3, AMPK, phospho-AMPK, PDH, phospho-PDH, MFN2, OGT, GFAP and loading control across genotype and age. n = 3 mice per group. **(E-J)** Quantification of NOTCH3 extracellular-domain signal **(E)**, phospho-AMPK/AMPK **(F)**, phospho-PDH/PDH **(G)**, MFN2 **(H)**, OGT **(I)** and GFAP **(J)**. **(K)** Representative hippocampal CA2 NOTCH3 extracellular-domain (green) and TOM20 (red) and merge with DAPI (blue) staining at six months. **(L-N)** NOTCH3-TOM20 spatial association **(L)**, relative TOM20 fluorescence intensity **(M)** and TOM20-positive volume **(N)**. Scale bar = 50μm. n = 3 mice per group, 5 fields per mouse were used for quantification. Data are presented as mean ± SEM, *p < 0.05, **p < 0.01 by Welch’s t-test.

Bulk cortical RNA sequencing revealed distinct transcriptional changes at three and six months. At three months, ranked KEGG analysis identified negative enrichment of glycolysis/gluconeogenesis and carbon metabolism, alongside positive enrichment of oxidative phosphorylation (**Fig. 6B**). These changes were accompanied by reduced expression of *Ldhd* (log₂FC = -0.48), a gene involved in lactate metabolism. By six months, carbon metabolism and the TCA cycle were negatively enriched, suggesting broader alterations in central carbon metabolic pathways (**Fig. 6C**). Increased expression of *Car13* (log₂FC = 1.18), involved in carbon dioxide and bicarbonate handling, was also observed. Immune-related pathways, including antigen processing and presentation, were more prominently enriched at six months, indicating differences in metabolic and inflammatory transcriptional programmes across the studied ages.

Further analyses revealed changes in interferon and TNFα/NF-κB signalling, together with extracellular matrix organisation and complement-related pathways (**Supplementary Fig. 7A-C**). Notably, several complement components and regulatory genes showed increased expression at six months, consistent with the complement-associated transcriptional changes identified in astrocytes from human MT BBB-like microtissues (**Fig. 3F**). The findings suggest that complement and inflammatory signalling become more prominent in the older NOTCH3^KI^ brain, alongside persistent alterations in metabolic and vascular-associated pathways.

Western blotting of mouse cortical brain tissue was used to assess whether the transcriptomic observations were accompanied by protein-level differences. Representative blots included NOTCH3, AMPK and phosphorylated AMPK, PDH and phosphorylated PDH, MFN2, OGT and GFAP across genotypes and ages (**Fig. 6D**). Quantification revealed increased NOTCH3 extracellular-domain signal in NOTCH3^KI^ mice at both ages. MFN2, OGT and GFAP were also higher in NOTCH3^KI^ samples, while phosphorylation-associated changes differed between ages. The phospho-PDH/PDH ratio was lower in NOTCH3^KI^ mice at three months, whereas phospho-AMPK/AMPK was reduced at six months (**Fig. 6D-J**).

Immunofluorescence at six months further showed NOTCH3 and TOM20 staining in the sampled brain regions (**Fig. 6K**). Six-month NOTCH3^KI^ group exhibited higher TOM20 fluorescence intensity, TOM20-positive volume and spatial association with NOTCH3 staining. (**Fig. 6L-N**).

Thus, the mouse data place the cellular findings within a temporal disease framework: metabolic dysregulation emerges early and precedes broader neurovascular, matrix, and inflammatory remodelling. We next tested this progression *in vivo* using complementary metabolic and vascular imaging.

### Multimodal imaging validates early metabolic dysfunction and progressive vascular remodelling in NOTCH3 p.R544C mice

We next combined metabolic and vascular imaging to determine whether the molecular differences in the NOTCH3^KI^ mouse model were accompanied by measurable changes in brain physiology and vessel architecture^78^. Structural MRI did not reveal overt morphological abnormalities in three-month-old NOTCH3^KI^ mice compared with WT controls (**Supplementary Fig. 8**). Despite the absence of apparent gross structureal changes at three months, hyperpolarised [1-¹³C]pyruvate magnetic resonance measurements were used to assess pyruvate-derived cerebral signals (**Fig. 7A,B**). NOTCH3^KI^ mice exhibited higher bicarbonate-to-pyruvate and alanine-to-pyruvate ratios than WT controls (**Fig. 7C,D**). Consistent with our transcriptomic analysis, these findings identify altered pyruvate metabolism as a potential early metabolic phenotype of NOTCH3^KI^ mice.

**Figure 7.**
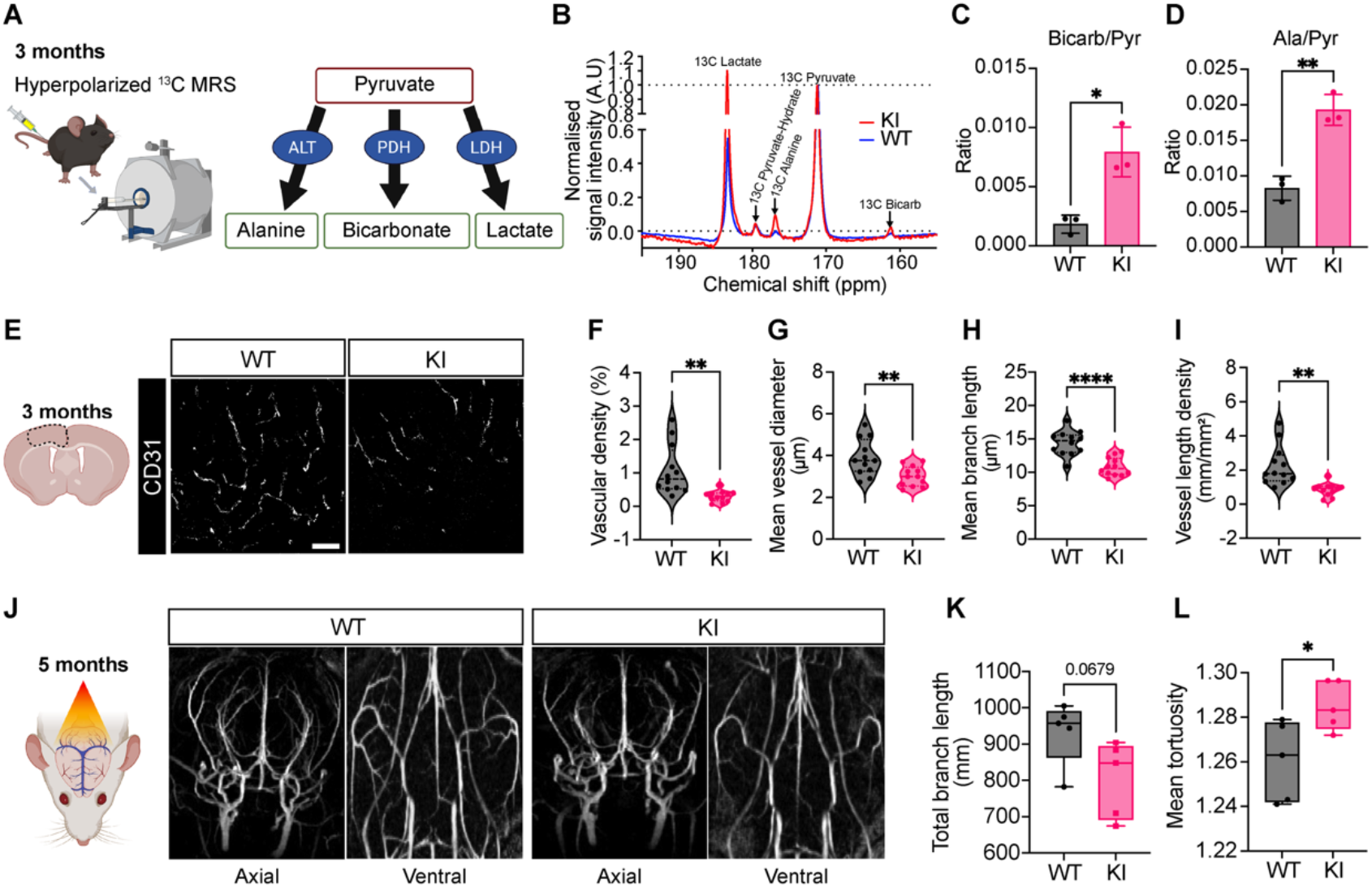
NOTCH3 p.R544C mice show altered cerebral pyruvate metabolism and vascular architecture. **(A)** Schematic of hyperpolarised [1-^13^C]pyruvate imaging in three-month-old NOTCH3^KI^ vs WT mice and major pyruvate-derived products. **(B)** Representative cerebral hyperpolarised ^13^C spectra. **(C,D)** Bicarbonate-to-pyruvate **(C)** and alanine-to-pyruvate **(D)** ratios. **(E)** Representative CD31-labelled cortical microvessels at three months. **(F-I)** Quantification of vessel density **(F)**, mean vessel diameter **(G)**, mean branch length **(H)** and vessel-length density **(I),** 5-7 fields across 3 mice per group were used. **(J)** Representative 3D vascular images and corresponding processed views at five months. **(K,L)** Quantification of total branch length **(K)** and mean vessel tortuosity **(L)**. Data are presented as mean ± SEM, *p < 0.05, **p < 0.01, ****p < 0.0001 by Welch’s t-test.

Microvascular abnormalities were also evident at three months. Representative CD31-labelled cortical vessels suggested differences in vascular organisation (**Fig. 7E**), and quantitative analysis showed lower vessel density, mean vessel diameter, mean branch length and vessel-length density in NOTCH3^KI^ than WT mice (**Fig. 7F-I**). Thus, altered pyruvate-derived signals and microvascular changes were both detectable at this age.

At five months, 3D vascular reconstructions using magnetic resonance angiography enabled assessment of a broader vascular field (**Fig. 7J**). Compared with WT controls, NOTCH3^KI^ mice exhibited modestly reduced total branch length and increased mean vessel tortuosity in the Circle of Willis (**Fig. 7K,L**). Consistent with the CD31 immunolabelling findings, these results support early vascular abnormalities in NOTCH3^KI^ mice. We conducted behavioural studies to determine whether the NOTCH3^KI^ mice showed early deficits. Behavioural assessment at three months revealed reduced time spent and distance travelled in the central zone during the open-field test, together with reduced immobility in the tail-suspension test. NOTCH3^KI^ mice also exhibited increased spontaneous alternation in the Y-maze without significant changes in total arm entries. These findings suggest altered exploratory and anxiety-related behaviour, without consistent evidence of cognitive impairment across the behavioural tests performed (**Supplementary Fig. 9**).

Together, these multimodal data define a temporal sequence in which altered cerebral metabolism and microvascular structure emerge early, followed by increasing mitochondrial and vascular remodelling with age. Integrated with the human BBB and single-cell findings, these results support an early multicellular neurovascular disease state that precedes more advanced structural injury and memory loss.

## Discussion

Our findings define NOTCH3 p.R544C CADASIL as an early multicellular neurovascular disorder organised across three interconnected mechanistic layers: BBB failure, complement activation, and metabolic dysregulation. First, MT NOTCH3 establishes an intrinsic endothelial disease state characterised by impaired angiogenesis, junctional disorganisation, and reduced barrier function, which persists in multicellular BBB models and develops into maladaptive vascular remodelling. Second, single-cell profiling reveals that endothelial BBB programme loss is accompanied by astrocytic complement, innate immune, and extracellular-matrix activation, indicating that the vascular defect propagates into a broader multicellular inflammatory state. Third, impaired mitochondrial respiratory capacity, altered cerebral pyruvate metabolism, and age-dependent metabolic changes identify metabolic dysregulation as an additional early disease layer that precedes more extensive neurovascular and inflammatory remodelling in vivo.

Importantly, these processes do not appear to represent independent abnormalities. Rather, the convergence of endothelial barrier failure, complement-matrix signalling, and altered metabolism suggests a coordinated early neurovascular disease programme that emerges before advanced structural brain injury. The improvement in BBB function and attenuation of selected complement-matrix signals following cilostazol and ISMN treatment further indicate that components of this early state remain pharmacologically modifiable. Together, these findings provide a framework linking endothelial dysfunction with broader multicellular neurovascular abnormalities and age-associated disease phenotypes, while suggesting that BBB dysfunction may represent a therapeutically modifiable feature of early NOTCH3-associated small vessel disease.

### NOTCH3 p.R544C establishes an early endothelial and BBB disease state

This study identifies early multicellular neurovascular dysfunction as a central feature of NOTCH3 p.R544C CADASIL and extends the conventional mural-cell-centred view of disease pathogenesis. Using complementary human and *in vivo* models, we show that p.R544C is associated with an intrinsic endothelial phenotype characterised by impaired angiogenic remodelling, migration, junctional organisation and barrier function. These abnormalities persisted in a 3D multicellular environment, where MT microtissues formed patent vascular networks but exhibited enlarged luminal structures and markedly increased permeability. Although the enlarged lumen phenotype was not recapitulated in vivo, the reduced microvascular density and altered vessel architecture observed in NOTCH3^KI^ mice provide complementary evidence of vascular abnormalities across experimental models. This distinction is important: p.R544C does not abolish vascular formation, but instead permits the formation of structurally established yet functionally unstable microvessels.

Previous studies of CADASIL have largely focused on degeneration of vascular smooth muscle cells and pericytes, consistent with the high expression of NOTCH3 in mural cells and the accumulation of NOTCH3 extracellular domain and granular osmiophilic material within vessel walls. Our findings do not challenge the importance of mural-cell pathology, but broaden this framework by identifying endothelial and BBB dysfunction as additional early components of the disease process. The convergence of impaired endothelial migration, reduced transendothelial electrical resistance, discontinuous ZO1 organisation and dysregulation of angiogenic, focal-adhesion and extracellular-matrix pathways supports a coherent endothelial disease state rather than an isolated permeability phenotype.

The 3D BBB-like microtissue model extends the endothelial phenotype to a multicellular environment containing astrocytes and pericytes, revealing distinct transcriptional responses across neurovascular cell populations. Endothelial cells exhibited alterations in vascular development, cellular organisation and BBB-associated transport programmes, consistent with the impaired network formation, junctional organisation and increased permeability observed experimentally. Astrocytes, despite retaining relatively preserved morphology in monolayer culture, showed prominent changes in extracellular matrix and complement-associated programmes, including increased expression of C1R, C1S and CFH. These changes suggest that mutant astrocytes may contribute to an altered inflammatory and extracellular signalling environment surrounding the vasculature.

Pericytes exhibited changes in pathways associated with vasculature development, cell adhesion, chemotaxis and extracellular matrix organisation, raising the possibility that altered mural-cell support contributes to vascular instability and remodelling. Glial progenitor-like cells showed additional changes in morphogenesis, cell junction organisation and developmental pathways, indicating that the transcriptional effects of p.R544C extend beyond differentiated neurovascular populations. Although these findings implicate multiple cell types in the disease-associated response, their individual contributions to barrier permeability and vascular architecture remain unresolved. As the MT microtissues contain multiple cell populations carrying the variant, future studies using selective replacement of mutant and corrected cell types will be necessary to distinguish endothelial-intrinsic defects from the contributions of astrocytes and pericytes.

### Endothelial dysfunction is accompanied by an early astrocytic complement-matrix response

A major finding of this study is that endothelial dysfunction occurs together with a distinct astrocytic molecular response. Although MT astrocytes retained relatively preserved morphology under basal conditions, transcriptomic profiling demonstrated substantial changes in inflammatory and extracellular-matrix-associated programmes. This dissociation between preserved morphology and altered molecular state raises the possibility that astrocytic molecular dysfunction may occur before overt morphological reactivity.

Single-cell analysis of the multicellular microtissues further resolved these responses. Endothelial cells showed reduced expression of genes within the curated BBB-associated transport and efflux module, particularly ABCG2, alongside alterations in tight-junction-associated genes, including CLDN5, OCLN and TJP1. In contrast, astrocytes exhibited increased expression of complement-associated genes, including C1R, C1S, CFH and SERPING1, consistent with broader changes in complement and innate immune signalling.

Complement signalling provides a biologically plausible interface between vascular dysfunction, inflammation and extracellular-matrix remodelling. Locally produced complement components can influence inflammatory signalling, vascular integrity and matrix turnover, while extracellular-matrix changes can in turn modify endothelial adhesion, migration, vessel geometry and barrier stability. The identification of matrix-associated abnormalities in both endothelial cells and astrocytes therefore suggests that complement and extracellular-matrix regulation may represent interconnected features of p.R544C-associated neurovascular dysfunction.

Importantly, the present data establish pathway convergence but not directionality. We cannot determine whether endothelial dysfunction promotes astrocytic complement activation, whether complement contributes directly to barrier failure, or whether both arise in parallel downstream of NOTCH3-associated stress. Direct perturbation of complement components will therefore be required to establish whether this pathway is causal, compensatory or secondary.

### Complement and extracellular-matrix dysregulation converges across human models and patient plasma

The relevance of the complement-matrix programme was strengthened by convergence across independent human datasets. C1S contributed to the astrocytic complement signature and was also increased in CADASIL patient plasma, while CLU and VTN further linked complement regulation with extracellular-matrix biology. The MT microtissue secretome similarly showed changes in mediators involved in vascular remodelling and matrix regulation, including VEGF, uPA, TIMP-1, thrombospondin-1 and IGFBP-2.

These observations support the biological relevance of complement-matrix dysregulation beyond a single *in vitro* platform and provide translational support for pathways identified in the human cellular models and nominate specific mediators for targeted validation.

The convergence of endothelial, astrocytic, secreted and circulating signals is particularly relevant to CADASIL, in which NOTCH3 extracellular-domain accumulation is accompanied by abnormal deposition of extracellular-matrix proteins within the vessel wall. Our findings raise the possibility that altered matrix biology reflects not only passive protein accumulation but also an active remodelling response involving both vascular and glial compartments.

### Altered metabolic regulation is an early feature of NOTCH3 p.R544C-associated neurovascular disease

An important finding of this study is the presence of altered cerebral metabolic handling in young NOTCH3^KI^ mice. At three months, hyperpolarized [1-¹³C]pyruvate imaging revealed increased bicarbonate-to-pyruvate and alanine-to-pyruvate ratios, indicating altered partitioning of pyruvate-derived metabolic signals through PDH-associated decarboxylation and alanine transamination pathways. These findings identify an early functional metabolic phenotype associated with NOTCH3 p.R544C and extend the molecular characterisation of CADASIL beyond conventional vascular and histopathological measurements.

The accompanying molecular analyses provided further evidence of altered metabolic regulation. At three months, NOTCH3^KI^ mice exhibited reduced inhibitory PDH phosphorylation, consistent with the increased bicarbonate-to-pyruvate ratio detected by hyperpolarised ¹³C imaging, potentially reflecting altered regulation of PDH-mediated pyruvate oxidation. Increased MFN2 and OGT expression at both ages further suggested changes in mitochondrial dynamics and nutrient-sensitive metabolic regulation. By six months, reduced AMPK phosphorylation relative to WT mice, together with negative enrichment of TCA cycle and carbon metabolism pathways, indicated additional alterations in cerebral metabolic regulation. Although these findings involve different aspects of metabolism, they support the presence of metabolic abnormalities across the disease stages examined. However, direct measurements of PDH enzymatic activity and mitochondrial function in brain tissue will be required to establish the functional consequences of these molecular changes.

In parallel, mitochondrial respiration was altered in cells expressing MT NOTCH3, characterised by reduced basal and ATP-linked oxygen consumption, accompanied by changes in proton leak and maximal respiratory capacity. These findings provide functional evidence that NOTCH3 p.R544C is associated with altered mitochondrial bioenergetics. Notably, the increased bicarbonate-to-pyruvate ratio observed *in vivo*, together with the reduced respiratory capacity in the cellular model, suggests that NOTCH3 p.R544C may perturb mitochondrial metabolic regulation at multiple levels. While these observations reflect distinct aspects of cellular metabolism, they offer complementary insights into the metabolic consequences of MT NOTCH3 expression. Hyperpolarized ¹³C imaging captures dynamic, substrate-specific metabolic conversion in vivo, whereas extracellular flux analysis provides a broader assessment of mitochondrial respiratory function. Their convergence supports the presence of altered metabolic homeostasis associated with NOTCH3 p.R544C and highlights the value of combining functional metabolic imaging with cellular bioenergetic measurements to characterise early metabolic changes in CADASIL.

The metabolic phenotype was accompanied by microvascular abnormalities already detectable at three months, while six-month-old mice exhibited more prominent inflammatory, extracellular-matrix and gliovascular changes. As the age-stratified cohorts were examined cross-sectionally, these observations do not establish a direct temporal or causal relationship between metabolic dysfunction and subsequent vascular remodelling. Nevertheless, they suggest that altered cerebral metabolism is an early component of the NOTCH3 p.R544C-associated disease phenotype and may provide a functional readout of neurovascular disturbance.

The cellular origins of these metabolic changes remain unresolved. Altered endothelial substrate transport, cerebral perfusion, astrocytic metabolic support and mitochondrial regulation may all contribute to the observed phenotype. Future studies combining cell-resolved metabolic profiling with longitudinal hyperpolarized imaging and vascular assessment will be important for determining how metabolic abnormalities relate to BBB dysfunction and progressive cerebrovascular pathology.

### Pharmacological rescue identifies a modifiable component of the early neurovascular phenotype

The improvement in barrier function following combined cilostazol and isosorbide mononitrate treatment provides an important functional dimension to the study. Treatment reduced permeability in MT microtissues and attenuated selected complement- and extracellular-matrix-associated proteins, including C1S and VTN, indicating that components of the disease-associated functional and molecular phenotype are pharmacologically modifiable.

These agents act through complementary cyclic nucleotide pathways and have been investigated in CSVD. The present experiments do not establish their precise mechanism of action in p.R544C CADASIL, nor do they demonstrate that suppression of complement-matrix signalling mediates improvement in barrier function. Nevertheless, the findings provide proof-of-principle that at least part of the early neurovascular phenotype is reversible in the human microtissue model.

This observation is particularly relevant in the context of a genetic disease. Although the underlying NOTCH3 mutation remains unchanged, downstream vascular abnormalities can nevertheless be modified experimentally. Targeting BBB instability, neurovascular signalling or associated inflammatory and matrix pathways may therefore provide an opportunity to intervene during an earlier phase of disease, before more advanced vessel and tissue injury becomes established.

## Summary

Taken together, our findings support a model in which NOTCH3 p.R544C establishes an intrinsic endothelial disease state characterised by impaired vascular remodelling, transport and barrier integrity. Within a multicellular neurovascular environment, this state is accompanied by astrocytic complement activation and extracellular-matrix remodelling and is associated with the formation of structurally patent but functionally unstable microvessels. *In vivo*, metabolic abnormalities are evident at three months, whereas vascular, extracellular-matrix, glial and inflammatory changes become more prominent by six months. This pattern suggests that early functional and vascular disturbances are accompanied by broader molecular changes at later disease stages, potentially contributing to progressive neurovascular dysfunction.

This model extends the traditional emphasis on mural-cell degeneration by placing BBB dysfunction and multicellular neurovascular signalling within an early phase of CADASIL pathogenesis. It also provides a framework linking molecular changes in human cellular models with circulating signals and age-associated abnormalities *in vivo*. Although the directionality between endothelial dysfunction, complement activation and metabolic disturbance remains to be established, their convergence across experimental systems supports their relevance to p.R544C disease biology.

### Study limitations and future directions

Several limitations should be considered together with opportunities for future investigation. The BBB-like microtissue captures key multicellular features of the neurovascular unit but does not fully reproduce physiological haemodynamic forces, systemic immune interactions, or the full cellular complexity of the adult brain. Future studies using in vivo models will help determine how these factors influence the neurovascular phenotype.

The mechanistic relationships among BBB failure, complement activation, and metabolic dysregulation also remain incompletely resolved. Although single-cell profiling identifies coordinated cell-specific programmes, it does not establish causal directionality. Direct perturbation of complement signalling, together with targeted manipulation of endothelial and astrocytic pathways, will be important to determine whether complement activation drives BBB dysfunction or represents a secondary response.

The cellular origin of the early metabolic phenotype also remains uncertain because bulk RNA sequencing lacks cell-type resolution and hyperpolarised ^13^C imaging reflects integrated tissue metabolism. Spatial transcriptomic, single-cell, and cell-specific metabolic approaches could therefore be used to identify the cellular and regional sources of altered pyruvate handling and link them directly to vascular and inflammatory changes.

Finally, longitudinal intervention studies integrating metabolism, perfusion, BBB permeability, and three-dimensional vascular imaging will be important to determine whether targeting these early neurovascular abnormalities alters subsequent disease progression. Such studies should help establish the temporal and causal relationships among BBB failure, complement activation, and metabolic dysregulation, and define which components represent the most tractable therapeutic targets.

## Conclusion

Our findings identify NOTCH3 p.R544C CADASIL as an early disorder of the multicellular neurovascular unit. The mutation is associated with intrinsic endothelial dysfunction, BBB instability, astrocytic complement activation, extracellular-matrix remodelling and early metabolic abnormalities, with broader neurovascular and inflammatory changes becoming more prominent at later ages.

By linking these processes across isogenic human models, 3D BBB-like microtissues, patient plasma and age-stratified mouse studies, this work broadens the mechanistic framework of CADASIL beyond mural-cell degeneration alone. Importantly, pharmacological improvement of barrier function demonstrates that components of this early neurovascular phenotype remain modifiable, identifying a potential disease window in which BBB-centred dysfunction could be targeted before more advanced cerebrovascular injury develops.

## Materials and Methods

### Human samples and ethics

Human blood samples were collected from patients recruited at the National Neuroscience Institute, Singapore (NNI), following written informed consent. All procedures were conducted in accordance with institutional guidelines and relevant regulations. Clinical diagnosis and pathological assessment were performed by clinicians and pathologists at NNI. The study was approved by the NNI Institutional Review Board (2017/2602).

### Generation and maintenance of human iPSCs

Peripheral blood mononuclear cells isolated from patients with CADASIL carrying the NOTCH3 p.R544C variant were reprogrammed into induced pluripotent stem cells (iPSCs) using the CytoTune-iPSC Sendai Reprogramming Kit (Thermo Fisher Scientific, A16517) according to the manufacturer’s instructions. Sendai reprogramming vectors were removed by subcloning iPSC clones over five passages. Following five passages, RNA was isolated using the PrimeWay Total RNA extraction kit (1st Base, KIT-9021) and reverse-transcribed using the High Capacity cDNA Reverse Transcription Kit (Applied Biosystems, 4368814). Clearance of Sendai vectors was confirmed by PCR using the primers 5′-GGATCACTAGGTGATATCGAGC-3′ and 5′-ACCAGACAAGAGTTTAAGAGATATGTATC-3′. Confirmed iPSC clones were cryopreserved in CryoStor CS10 freezing medium (BioLife Solutions, 210102). Cells were maintained on Matrigel-coated culture vessels (Corning, 354277) in complete mTeSR PLUS medium (STEMCELL Technologies, 85850) at 37 °C and 5% CO₂ and passaged weekly using ReLeSR dissociation reagent (STEMCELL Technologies, 100-0483).

### Generation of the isogenic corrected control

NOTCH3 p.R544C iPSCs were expanded to approximately 70% confluence and dissociated using TrypLE to obtain a single-cell suspension. Cell viability was determined by trypan blue exclusion. Ribonucleoprotein complexes were generated by incubating 37.5 pmol sgRNA (5′-CACGCTGTGTGATTGCAACG-3′; Integrated DNA Technologies), 1 μg spCas9 protein (Integrated DNA Technologies) and 1.25 μl R buffer (Neon Transfection Kit, Invitrogen, MPK10096) for 20 min at room temperature. A single-stranded oligodeoxynucleotide donor template (0.5 μg) was subsequently added. Ribonucleoprotein complexes were mixed with 2 × 10⁵ iPSCs and electroporated using the Neon Transfection System at 1050 V, 30 ms and two pulses. Electroporated cells were seeded at 500-5000 cells cm⁻². Clonal populations were expanded and genomic DNA was extracted using QuickExtract DNA Extraction Solution (Lucigen, QE09050). Clones were genotyped by PCR using the Q5 High-Fidelity PCR Kit (New England Biolabs, M0491L).

### Differentiation of iPSCs into endothelial cells and pericytes

Endothelial cells and pericytes were differentiated from iPSCs using a lateral mesoderm induction protocol as previously described^79,80^. Growth factor-reduced Matrigel (Corning, 356231) was diluted 1:30 and used to coat culture vessels for 1 h at room temperature. Immediately before seeding, Matrigel was removed and plates were washed once with PBS. Pre-warmed mTeSR PLUS supplemented with 10 μM Y-27632 (Cellagen, C9217-10) was added. iPSCs were dissociated using Accutase (Gibco, A1105-01), washed and seeded at 47,000 cells cm⁻². For lateral mesoderm induction, cells were cultured from days 1-3 in N2B27 medium supplemented with 7 μM CHIR-99021 (STEMCELL Technologies, 100-1042) and 25 ng ml⁻¹ human BMP4 (STEMCELL Technologies, 78211), without medium replacement. On days 4-5, endothelial induction was performed using StemPro-34 SFM (Gibco, 10639011) supplemented with 200 ng ml⁻¹ VEGF165 (PeproTech, 100-20) and 2 μM forskolin (Abcam, ab120058), with daily medium replacement. On day 6, CD144-positive ECs were enriched by magnetic-activated cell sorting (MACS; Miltenyi Biotec, 130-097-857) and replated onto fibronectin-coated vessels. The CD144-negative fraction was plated separately on vessels coated with Cell Systems Attachment Factor and maintained in Cell Systems culture medium (CSS-A101) to support pericyte outgrowth. From day 7 onwards, ECs were expanded in EGM-MV2 medium, while pericytes were maintained using the Cell Systems culture kit. Three independent differentiations were performed for the main endothelial cell, astrocyte and pericyte experiments. Cellular identity was confirmed using immunofluorescence staining of cell-type specific markers (**Supplementary Fig. 1**)

### Differentiation of iPSCs into astrocytes

Astrocytes were generated from mature cerebral organoids using the STEMdiff Cerebral Organoid Kit (STEMCELL Technologies, 8570)^81^. iPSC colonies were dissociated into single cells using Accutase and 8,000 cells were seeded per well in V-bottom ultra-low-attachment 96-well plates (Sbio, MS-9096VZ) in embryoid body formation medium supplemented with 10 μM Y-27632. Medium was refreshed on days 2 and 4. On day 5, embryoid bodies were transferred to 24-well low-attachment plates and cultured in neural induction medium for 2 days. Each embryoid body was subsequently embedded in Matrigel for Organoid Culture (Corning, 356255) and cultured in neural expansion medium for a further 3 days. Cerebral organoids were then transferred to maturation medium and maintained on an orbital shaker, with medium replacement every 3 days. Astrocytes were isolated from day-90 cerebral organoids using enzymatic dissociation followed by magnetic enrichment. Five organoids were mechanically disrupted and digested with papain (Worthington Biochemical, LK003150) to obtain a single-cell suspension. Cells were plated onto Matrigel-coated vessels in astrocyte culture medium (ScienCell, 1801) and passaged at least twice before enrichment. Astrocytes were purified by MACS using an anti-GLAST antibody (Miltenyi Biotec, 130-096-052), seeded at 15,000 cells cm⁻² and maintained in astrocyte culture medium.

### Astrocyte morphology analysis

iPSC-derived astrocytes were immunostained for GFAP and counterstained with 4’6-diamidino-2-phenylindole dihydrochloride (DAPI) (Sigma-Aldrich, #B2261). Confocal images were acquired under identical settings across groups. Astrocyte morphological complexity was quantified by Sholl analysis using the Neuroanatomy plugin in Fiji/ImageJ. Individual GFAP-positive astrocytes were traced and process branching was quantified as a function of radial distance from the cell soma.

### Endothelial angiogenesis assay

Angiogenic network formation was assessed using µ-Slide 15-well 3D plates (ibidi, 81501). Growth factor-reduced basement membrane matrix (Corning, 354230) was prepared according to the manufacturer’s instructions and 10 μl was added to the inner well of each chamber. Plates were incubated at 37 °C for 1 h to allow polymerisation. Endothelial cells were dissociated using TrypLE, resuspended at 1 × 10⁵ cells ml⁻¹ and 50 μl of cell suspension was added to each well. Angiogenic networks were imaged using an inverted light microscope. Network parameters were quantified using AngioTool2/Fiji and included vessel length, vessel diameter, number of junctions and vessel area. Ten wells were analysed per line.

### Endothelial migration assay

Cell migration was assessed using a scratch wound-healing assay. ECs were seeded at matched densities and grown to near confluence. A linear scratch was introduced using a sterile pipette tip and detached cells were removed by washing with PBS. Fresh medium was added and cultures were imaged immediately and after 18 h. Migration was quantified from the remaining uncovered area relative to the initial wound area using Fiji/ImageJ.

### Transendothelial electrical resistance

ECs were seeded onto 24-well Transwell inserts with a surface area of 0.33 cm² at 70,000 cells per insert in EGM-2 supplemented with 10% FBS. The apical and basolateral compartments contained 200 μl and 700 μl medium, respectively. After overnight attachment, medium was replaced with EGM-2 containing 2% heat-inactivated FBS. Cells were maintained in starvation medium for 4 h before the first measurement. Transendothelial electrical resistance was measured using an EVOM2 epithelial volt-ohm meter equipped with STX electrodes (World Precision Instruments). Measurements were subsequently acquired at 24-h intervals for a total of three readings. Medium was refreshed before each measurement. Blank insert resistance was subtracted and values were normalised to membrane surface area and expressed as Ω·cm².

### Cell line culture and NOTCH3 overexpression

HeLa-mKeima and HEK293T cells were maintained in high-glucose DMEM without glutamine supplemented with 10% fetal bovine serum (Gibco, A5256801), 1% penicillin-streptomycin (Gibco, 15140122), 1% non-essential amino acids (Gibco, 11140-050) and 1% GlutaMAX (Gibco, 35050-038). HeLa cells were passaged using TrypLE (Gibco, 12604013), while mKeima-HeLa cells were passaged using 0.25% trypsin (Gibco, 25200056). WT or MT NOTCH3 constructs were transfected using Lipofectamine LTX (Invitrogen, 15338100) according to the manufacturer’s instructions. Cells were harvested or fixed 48 h after transfection for western blotting or immunofluorescence, respectively.

### Mitophagy assay

mKeima-HeLa cells were transfected with wild-type or p.R544C NOTCH3 extracellular-domain constructs. At 48 h after transfection, cells were treated with vehicle (DMSO) or 30 μM FCCP for 4 h and imaged live by confocal microscopy. Images were analysed using QuPath.

### Seahorse mitochondrial stress test

Mitochondrial respiration and glycolytic rates were evaluated using a Seahorse XFe24 Extracellular Flux Analyzer (Agilent Technologies, USA). Transfected HEK293T cells were seeded at a density of 8 × 10^4^ cells/well on poly-D-lysine-coated Seahorse XF 24-well microplates. 1 hr before the measurement, the culture medium was replaced with XF minimal medium (supplemented with 10 mM glucose, 2 mM pyruvate, 4 mM glutamine). The oxygen consumption rate (OCR) and extracellular acidification rate (ECAR) were recorded according to the manufacturer’s instructions. For mitochondrial stress and glycolytic profiling, compounds were injected at the following final concentrations: 4 *μ*M oligomycin, 0.5 *μ*M FCCP, 2 *μ*M rotenone, and 2 *μ*M antimycin A.

### Generation of 3D BBB-like microtissues

BBB-like microtissues were generated using the OrganiX platform (AIM Biotech, AIMOGNXP). Fibrinogen (Sigma-Aldrich, T7513) was prepared at 6 mg ml⁻¹ and maintained on ice. Thrombin (Sigma-Aldrich, F8630) was diluted in EGM-MV2 supplemented with 50 ng ml⁻¹ VEGF and AGS to a final concentration of 1 U ml⁻¹. iPSC-derived endothelial cells, astrocytes and pericytes were combined at a ratio of 2:0.3:0.3. Cells were resuspended in chilled thrombin-containing medium. Equal volumes of thrombin-cell suspension and fibrinogen were mixed on ice and immediately loaded into the central chamber of the OrganiX device. Hydrogels were allowed to polymerise for 45 min at room temperature. Medium was subsequently introduced into the inlet channel and reservoirs were filled with unequal volumes to establish volume-driven interstitial flow. Medium was refreshed daily and reservoir volumes were alternated to maintain flow.

### Pharmacological treatment of BBB-like microtissues

Microtissues were treated with vehicle or a combination of 1 μM cilostazol (MedChemExpress, HY-17464) and 10 μM isosorbide mononitrate (MedChemExpress, HY-B0642). Treatment-containing medium was refreshed daily for 7 days.

### Dextran permeability assay

Vascular permeability was assessed 6 days after seeding using fluorescent dextran (Invitrogen, D34681). Dextran was introduced into the media channels and flow was generated by loading unequal volumes into opposing channels. Microtissues were inspected after 5-10 min to confirm uniform vascular perfusion before equilibration of channel volumes. Confocal z-stacks were acquired at multiple positions within each microtissue to account for spatial heterogeneity. Permeability was quantified using a permeability index calculated as fluorescence intensity outside the vessel lumen divided by fluorescence intensity within the vessel lumen.

### Microtissue lumen analysis

Confocal z-stacks of CD31-labelled vascular structures were analysed using Fiji/ImageJ or QuPath. Vascular lumen size was quantified from segmented CD31-positive structures. Five microtissues were analysed per condition, comprising 77 corrected and 66 MT luminal regions in the dataset used for the main analysis.

### Collection of microtissue-conditioned medium

Microtissue culture supernatants were collected and centrifuged at 10,000 × g for 10 min to remove cellular debris. Clarified supernatants were stored at −80 °C until analysis.

### Recovery of cells from microtissues

Microtissues were extracted from the microfluidic devices using the supplied plunger. Nattokinase (MedChemExpress, HY-P2373) was prepared at 2 mg ml⁻¹ in PBS and pre-warmed to 37 °C. Microtissues were submerged in 100 μl nattokinase solution and incubated for 30 min at 37 °C. Samples were washed in serum-free medium and pelleted by centrifugation at 300 × g for 10 min before downstream processing.

### Immunofluorescence staining and confocal imaging

Cells and microtissues were fixed in 4% paraformaldehyde for 20 min and 1 h, respectively, at room temperature. Mouse brain specimens were fixed with 4% paraformaldehyde and then soaked in 30% sucrose before use. The fixed specimens were then sliced by a cryostat machine (Leica). Samples were washed three times with PBS, permeabilised using 0.05% Triton X-100 and blocked in 1% bovine serum albumin in PBS. Primary antibodies were diluted in 1% BSA/PBS and incubated overnight at 4 °C. Following three washes, samples were incubated with fluorescent secondary antibodies overnight at 4 °C. DAPI (Sigma-Aldrich, #B2261) was used for nuclear counterstaining. Coverslips were mounted using FluorSafe mounting reagent (Millipore, 345789). Microtissues were imaged directly within the OrganiX plate. Mouse brain slices were mounted onto glass slides after staining. Images were acquired using confocal microscopy (FV3000, Olympus) and analysed using Fiji/ImageJ. Primary antibodies included NOTCH3 extracellular domain (1E4; Sigma-Aldrich, MABC594), CD31 (Cell Signaling Technology, 3528), PDGFRβ (Invitrogen, MA5-15143), GFAP (Novus Biologicals, NBP1-05198), CD144 (Invitrogen, 14-1449-82), ZO1 (Cell Signaling Technology, 13663S), SOX9 (Cell Signaling Technology, 82630).

### Western blotting

Brain tissue were lysed in RIPA buffer containing protease and phosphatase inhibitors. Lysates were maintained on ice for at least 30 min with intermittent vortexing and clarified by centrifugation at 13,000 rpm for 15 min at 4 °C. Protein concentration was determined using the DC Protein Assay (Bio-Rad, 5000112). Samples were mixed with loading buffer and denatured at 95 °C for 5 min. Equal amounts of protein (15 μg) were separated by SDS-PAGE using a Mini-PROTEAN system (Bio-Rad). Multi-percentage 4-8-15% acrylamide gels were used to resolve proteins across a broad molecular-weight range. Proteins were transferred to 0.45 μm PVDF membranes and blocked in 3% BSA in TBST for 1 h at room temperature.

Membranes were incubated with primary antibodies overnight at 4 °C followed by HRP-conjugated secondary antibodies for 2 h. Chemiluminescent signals were developed using ECL substrates (Thermo Fisher Scientific, 32209 and 34096) and acquired using a ChemiDoc MP system (Bio-Rad). Band intensity was quantified using Fiji/ImageJ. Protein expression was assessed for NOTCH3(1E4) (Sigma-Aldrich, MABC594), AMPK (Cell Signaling Technology, 2532), phospho-AMPK (Cell Signaling Technology, 2535), PDH (Cell Signaling Technology, 2784S), phospho-PDH (Cell Signaling Technology, 31866S), MFN2 (Cell Signaling Technology, 9482), OGT (Cell Signaling Technology, 24083) and GFAP (Chemicon, MAB360), with β-actin (CST, #3700) used as the loading control.

### Angiogenesis and cytokine proteome arrays

Conditioned medium from three microtissues per condition was pooled at Day 7. Microtissue secretomes were profiled using Human Angiogenesis Proteome Arrays (R&D Systems, ARY007) and Human XL Cytokine Arrays (R&D Systems, ARY022B) according to the manufacturer’s instructions. Culture supernatants were incubated with nitrocellulose membranes containing immobilised capture antibodies. Following overnight incubation, membranes were washed and incubated with biotinylated detection-antibody cocktails followed by streptavidin-HRP. Chemiluminescent signals were acquired using a ChemiDoc MP system and spot intensity was quantified after background subtraction using Fiji/ImageJ. Relative protein abundance was compared between experimental conditions.

### ELISA

CLU, VTN and C1S concentrations in microtissue-conditioned media and human plasma were measured using commercially available ELISA kits according to the manufacturers’ instructions. CLU was quantified using the Human Clusterin ELISA Kit (Invitrogen, EHCLU), while VTN and C1S were measured using the Human Vitronectin ELISA Kit (Novus Biologicals, NBP3-07095) and Human C1S ELISA Kit (Novus Biologicals, NBP3-27273), respectively. Samples were diluted 1:2 for CLU and C1S and 1:3 for VTN. Standards and diluted samples were added to pre-coated wells, followed by incubation with the respective biotinylated detection antibodies and streptavidin-HRP or HRP-conjugated detection reagents. Following substrate development with 3,3′,5,5′-tetramethylbenzidine (TMB), reactions were terminated using stop solution and absorbance was measured at 450 nm. Protein concentrations were calculated using four-parameter logistic standard curves. All standards and samples were analysed in duplicate.

### Single-cell RNA sequencing

Single-cell RNA-sequencing libraries were generated using the Chromium platform with Fixed RNA Profiling/Flex chemistry (10x Genomics). Two NOTCH3 p.R544C patient lines (120 and 121) and one isogenic corrected line (C51) were included. Twelve microtissue samples were analysed across three multiplexed plexes. Raw sequencing data were processed using Cell Ranger v10.0.0. Demultiplexing, alignment to GRCh38-2024-A, barcode assignment and unique molecular identifier counting were performed using the cellranger multi pipeline with probe-set-specific configuration files. Filtered feature-barcode matrices were imported into R v4.5.1 and analysed using Seurat v5.4.0. Sample metadata included sample identifier, patient identifier, experimental condition and plex assignment.

### Single-cell RNA-seq quality control and integration

Cells with fewer than 200 detected genes, more than 6,000 detected genes or >10% mitochondrial transcripts were excluded. Following filtering, datasets from the three plexes were integrated to reduce batch-associated variation. RNA layers were joined and SCTransform normalisation was performed with regression of mitochondrial transcript percentage. Integration features (n = 3,000) were selected using SelectIntegrationFeatures, followed by PrepSCTIntegration, FindIntegrationAnchors and IntegrateData. Principal component analysis was performed on the integrated dataset. Thirty principal components were used for UMAP embedding, neighbourhood graph construction and clustering. Initial clustering used a resolution of 0.4. Clusters with low feature counts, enrichment for ambient RNA-associated genes and strong sample-specific bias were excluded as technical artefacts. The cleaned dataset was subsequently reclustered using 30 principal components and a resolution of 0.2.

### Single-cell annotation and differential expression analysis

Cluster marker genes were identified using Seurat FindAllMarkers with min.pct = 0.25 and logfc.threshold = 0.25. Cell types were assigned based on canonical marker expression. Astrocytes were identified using markers including ALDH1L1, GFAP and SOX9; endothelial cells using CLDN5, OCLN, TJP1, MFSD2A, KDR, TEK, ABCG2 and SLC2A1; pericytes using PDGFRB, RGS5, MCAM and VTN; and glial progenitor-like cells using proliferative markers including MKI67, TOP2A and BUB1B. For disease-specific differential expression, each cell population was analysed separately using FindMarkers to compare CADASIL and corrected cells with min.pct = 0.1 and logfc.threshold = 0.25. Differential-expression results were combined across clusters for downstream pathway analysis using Metascape.

Selected complement components, complement regulators and extracellular matrix-associated genes were examined using previously generated single-cell differential expression results comparing MT and corrected BBB-like microtissues. Gene expression changes were extracted separately for endothelial cells, astrocytes, pericytes and glial progenitor-like cells and visualised using *ggplot2* in R. Bubble plots displayed log2 fold changes by colour and - log10(adjusted P value) by dot size. Genes with adjusted P < 0.05 and absolute log2 fold change ≥ 0.25 were indicated by asterisks. Genes absent from the exported differential expression dataset were marked with an ‘x’.

### NOTCH3 p.R544C mouse model generation

NOTCH3^KI^ mice were generated by Cyagen Biosciences using CRISPR/Cas9-mediated targeted insertion of MT human *NOTCH3* into the mouse *Rosa26* locus. Briefly, the knock-in expression cassette comprised the MT human *NOTCH3* coding sequence with C-terminal V5 and HA tags, flanked by mouse *Rosa26* homology arms. The gRNA sequence 5′-CTCCAGTCTTTCTAGAAGATGGG-3′ was used to target the *Rosa26* locus. The gRNA, CRISPR protein and donor DNA template were co-injected into fertilised mouse eggs to generate targeted offspring. Successful integration was confirmed by PCR amplification of the 5′ and 3′ homology-arm junctions, followed by Sanger sequencing. Heterozygous NOTCH3^KI^ mice were obtained and maintained for subsequent studies.

For genotyping, the knock-in allele was detected using primers 5′-TCCACCTACCCATACGATGTTC-3′ and 5′-ATACTCCGAGGCGGATCACAA-3′, generating a 503-bp product. The wild-type allele was detected using primers 5′-CACTTGCTCTCCCAAAGTCGCTC-3′ and 5′-ATACTCCGAGGCGGATCACAA-3′, generating a 453-bp product. WT mice lacking the knock-in allele served as controls. Mice were maintained in a specific-pathogen-free facility under a 12-h light/dark cycle, with food and water available ad libitum. Both male and female mice were included in the experiments. All animal procedures were conducted in accordance with institutional guidelines and approved by the Institutional Animal Care and Use Committee of the National Neuroscience Institute and Tan Tock Seng Hospital (A24007).

### Mouse brain bulk RNA sequencing

Total RNA was extracted from mouse brain cortical tissue using the miRNeasy Kit (Qiagen, 217084). mRNA sequencing was performed by Novogene using an Illumina HiSeq 4000 platform. Reads were aligned to the mouse reference genome (GRCm39), and gene-level read counts were generated using featureCounts v1.5.0-p3. Gene-expression levels were calculated as fragments per kilobase of transcript per million mapped reads. Differential expression analysis was performed separately for three- and six-month-old NOTCH3^KI^ and WT mice using DESeq2 in R. Genes were considered differentially expressed at a Benjamini-Hochberg-adjusted P value < 0.05. Ranked KEGG gene-set enrichment analysis (GSEA) was performed using DESeq2 Wald statistics and visualised as dot plots showing positively and negatively enriched pathways. For focused pathway analysis, preranked GSEA was performed using *fgsea* and gene sets from MSigDB, including GO Biological Process, KEGG, Reactome and Hallmark collections. Selected pathways were grouped into metabolism, vascular, extracellular matrix and inflammatory categories. Dot plots displayed normalised enrichment scores (NES), with dot size representing -log10 false discovery rate (FDR). Gene-level bubble plots were generated using selected genes that reached FDR < 0.05 at either age, with colour representing log2 fold change and dot size representing -log10(FDR). Complement-associated genes were additionally grouped into functional categories, including classical initiation, alternative amplification, central and terminal components, receptors and regulators. Gene-level changes were visualised using log2 fold changes and nominal P values, with asterisks indicating P < 0.05. All plots were generated using *ggplot2* in R.

### Hyperpolarised [1-¹³C]pyruvate magnetic resonance imaging

1-¹³C pyruvic acid (Cambridge Isotope Laboratories, US) containing 30 mM trityl radical (AH111501) was hyperpolarized using SpinAligner system (Polarize, Denmark) at an optimized microwave frequency. The polarized sample was dissoluted in 40mM of sodium dihydrogen phosphate buffer, pH 7.4 containing an equimolar concentration of NaOH, yielding an 80 mmol/L hyperpolarized 1-¹³C pyruvate solution at a physiological temperature and pH.

Animal experiments were performed on a 9.4T preclinical MRI system (Bruker, Germany). Mice were anesthetised with 4% isoflurane in an induction chamber and maintained under anaesthesia. A custom catheter was placed in the tail vein for intravenous hyperpolarised agent administration. The mouse was positioned prone on the scanner bed, and a dual-tuned 1H/¹³C surface coil was positioned over the head. Respiration rate and body temperature were continuously monitored throughout the experiment. Dynamic ¹³C MR spectra were acquired using the following parameters: repetition time TR = 2,000 ms, 60 repetitions, flip angle 10° and a central chemical shift set to 172.6 ppm.

Hyperpolarised [1-^13^C]pyruvate imaging was performed in 3-month-old mice. Cerebral bicarbonate-to-pyruvate and alanine-to-pyruvate ratios were calculated from integrated metabolite signals using MestReNova.

### 3D vascular imaging

3D time-of-flight (TOF) magnetic resonance angiography was performed using an 11.7 T Bruker BioSpec MRI system equipped with a 72-mm quadrature volume resonator for radiofrequency transmission and a four-element (2 × 2) cryogenic coil for signal reception. Mice were initially anaesthetised with 4% isoflurane and subsequently positioned on the MRI cradle, with the head secured using ear bars to minimise motion artefacts. Anaesthesia was maintained at 1.5% isoflurane in a medical air/oxygen mixture (0.6/0.3 L min⁻¹). Body temperature and respiratory rate were continuously monitored and maintained at 37 °C and above 100 breaths min⁻¹, respectively. Whole-brain angiographic images were acquired using a 3D TOF fast low-angle shot (3D-TOF-FLASH) sequence with generalised autocalibrating partially parallel acquisitions (GRAPPA). Acquisition parameters were as follows: field of view, 20 × 20 × 20 mm³; acquisition matrix, 360 × 360 × 360; echo time (TE), 2.0 ms; repetition time (TR), 12 ms; flip angle, 20°; and two signal averages. The total acquisition time was approximately 24 min. 3D TOF-MRA volumes were intensity-normalised and segmented using multiscale Hessian-based vessel enhancement with identical settings across scans^82,83^. Region of interest of the Circle of Willis was used for quantification. Mean vessel volume, length, branching, and tortuosity were quantified from the 3D masks and skeletons.

### Animal behavioural tests

Behavioural assessments were performed in three-month-old NOTCH3^KI^ and WT mice to evaluate locomotor activity, anxiety-related behaviour and learning and memory. The open-field test was used to quantify total distance travelled, exploratory activity and time spent in the central zone. Anxiety-related behaviour was further assessed using the elevated plus maze and elevated zero maze by measuring exploration of open and enclosed regions. Spontaneous alternation in the Y-maze was calculated from consecutive entries into all three arms, with total arm entries recorded as an additional measure of exploratory activity. Novel object recognition was assessed based on exploratory preference for novel versus familiar objects, while fear conditioning was evaluated using freezing-related measurements. Behavioural immobility was quantified using the tail-suspension test. Results were compared between NOTCH3^KI^ and WT mice, with individual animals considered biological replicates.

### Statistical analysis

Statistical analyses were performed using GraphPad Prism and R where appropriate. For comparisons between two groups, Welch’s unpaired t-test was used unless otherwise indicated. Comparisons involving more than two groups were performed using one-way analysis of variance followed by appropriate post hoc testing. Data are presented as mean ± SEM unless otherwise specified. A p < 0.05 was considered statistically significant. For transcriptomic analyses, multiple-testing correction was performed using the Benjamini-Hochberg method, with adjusted P < 0.05 considered significant unless otherwise stated. Exact sample sizes, definitions of n, statistical tests and measures of dispersion are provided in the corresponding figure legends.

## Data availability

Processed sequencing datasets and associated metadata will be made available upon reasonable request.

## List of Abbreviations

3D: Three-dimensional
3D-TOF-FLASH: 3D TOF fast low-angle shot
BBB: Blood-brain barrier
CADASIL: Cerebral autosomal dominant arteriopathy with subcortical infarcts and leukoencephalopathy
CSVD: Cerebral small vessel disease
DAPI: 4’6-diamidino-2-phenylindole dihydrochloride
ECAR: Extracellular acidification rate
FCCP: Carbonyl cyanide-trifluoromethoxyphenylhydrazone
FDR: False discovery rate
GRAPPA: Generalised autocalibrating partially parallel acquisitions
GSEA: Gene-set enrichment analysis
ISMN: Isosorbide mononitrate
MT: Mutant
NES: Normalised enrichment scores
NOTCH3KI: NOTCH3 knock-in
OCR: Oxygen consumption rate
TE: Echo time
TMB: 3,3’,5,5’-tetramethylbenzidine
TOF: Time-of-flight
TR: Repetition time
WT: Wild-type

