## Supplementary Figures for "NOTCH3 p.R544C Drives Early Neurovascular Unit Dysfunction through Blood-Brain Barrier Failure, Complement Signalling and Metabolic Dysregulation"

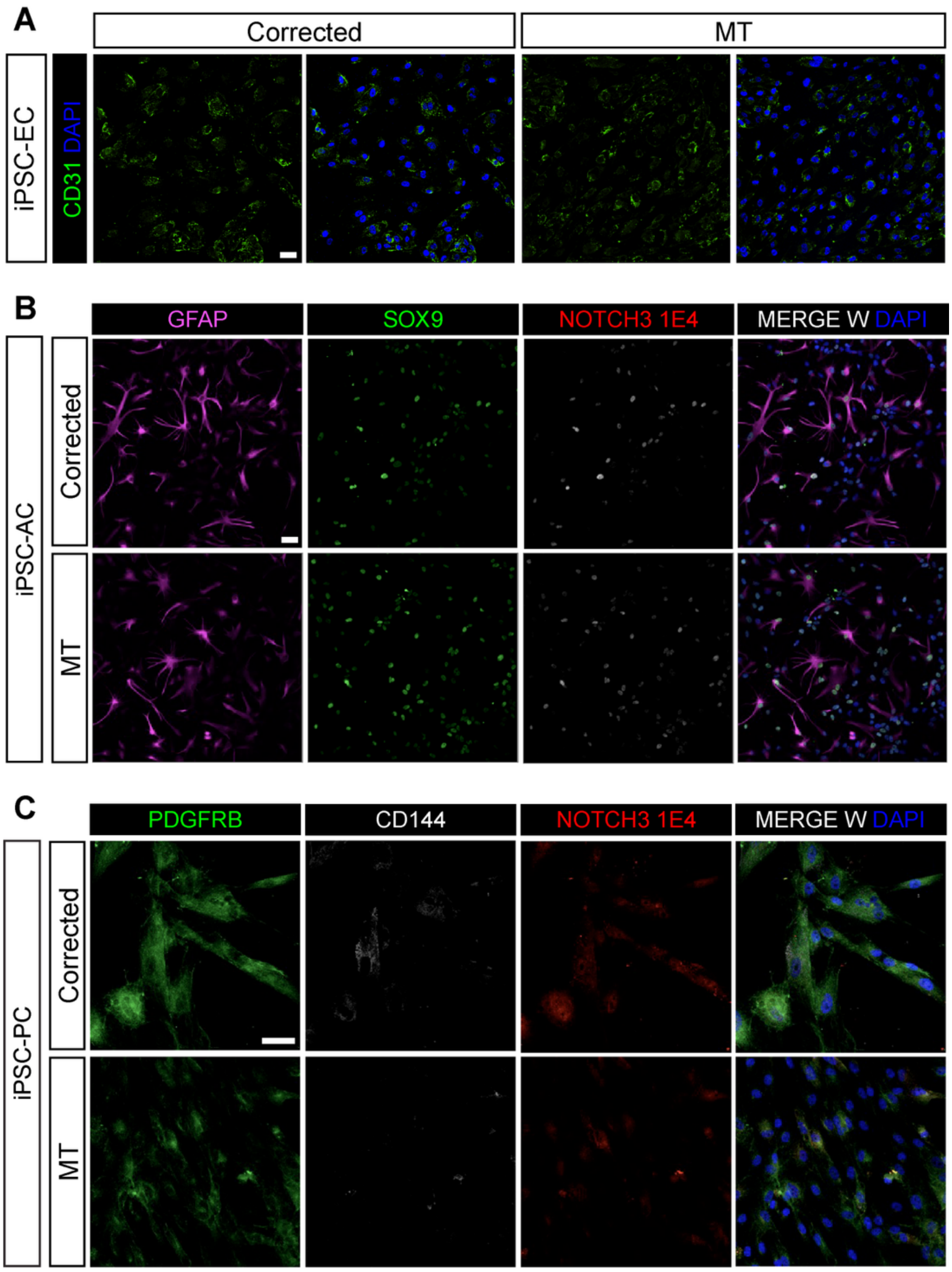


**Supplementary Figure 1. Characterisation of iPSC-derived endothelial cells, pericytes and astrocytes. (A)** Representative CD31 (green) merge with DAPI (blue) in corrected and MT ECs. **(B)** Representative GFAP (magenta), SOX9 (green) and NOTCH3 extracellular-domain (red) merge with DAPI (blue) staining in corrected and MT astrocytes. **(C)** Representative PDGFRβ (green), CD144 (white) and NOTCH3 extracellular-domain (red) merge with DAPI (blue) staining in corrected and MT pericytes. Scale bar = 50µm.

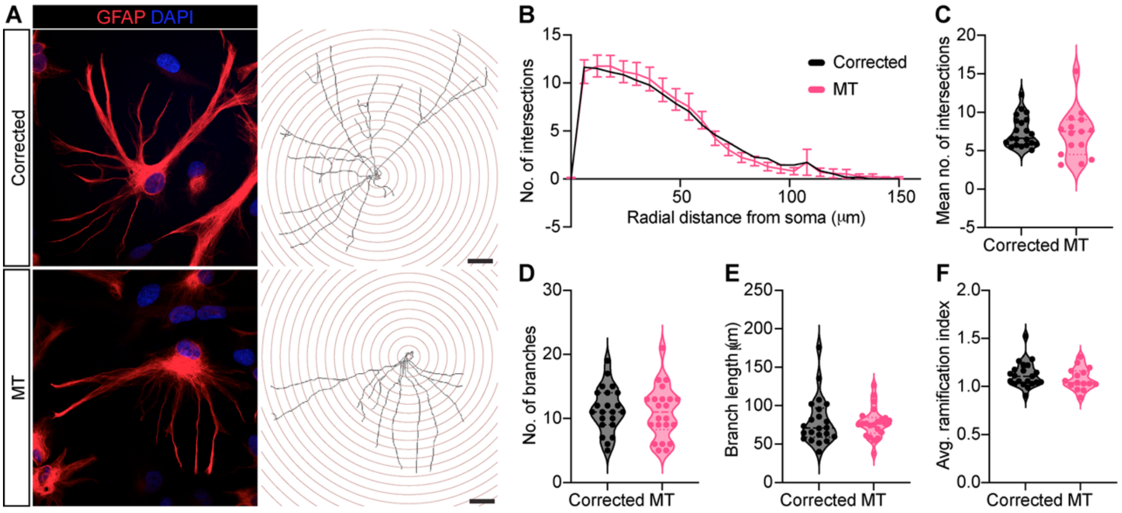


**Supplementary Figure 2. Morphological analysis of corrected and MT iPSC-derived astrocytes. (A)** Representative GFAP-positive astrocytes and corresponding Sholl reconstructions. **(B)** Sholl intersection profiles by radial distance from the soma. **(C-F)** Quantitative morphological parameters shown in the supplied supplementary figure, including process intersections and complexity-related measurements. Scale bar = 20μm. 22 cells per group across 3 batches were analysed. Data are presented as mean ± SEM, no significant difference by Welch's t-test.

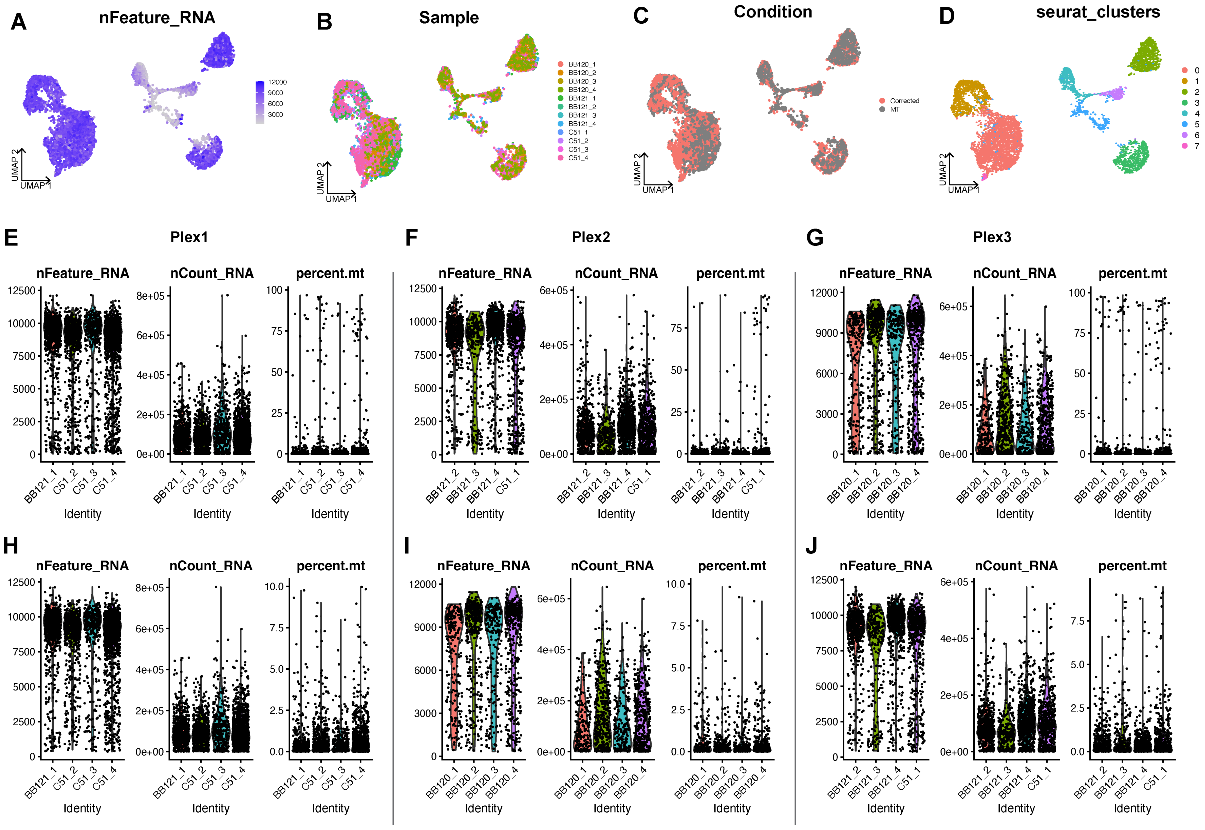


**Supplementary Figure 3. Integration of single-cell RNA-seq data. (A-D)** UMAP visualisations coloured by nFeature_RNA **(A)**, sample **(B)**, condition **(C)**, Seurat clusters before the exclusion of selected clusters due to low nFeature_RNA **(D)**. **(E-J)** Per-plex and per-sample quality-control distributions for detected genes, molecular counts and mitochondrial transcript percentages.

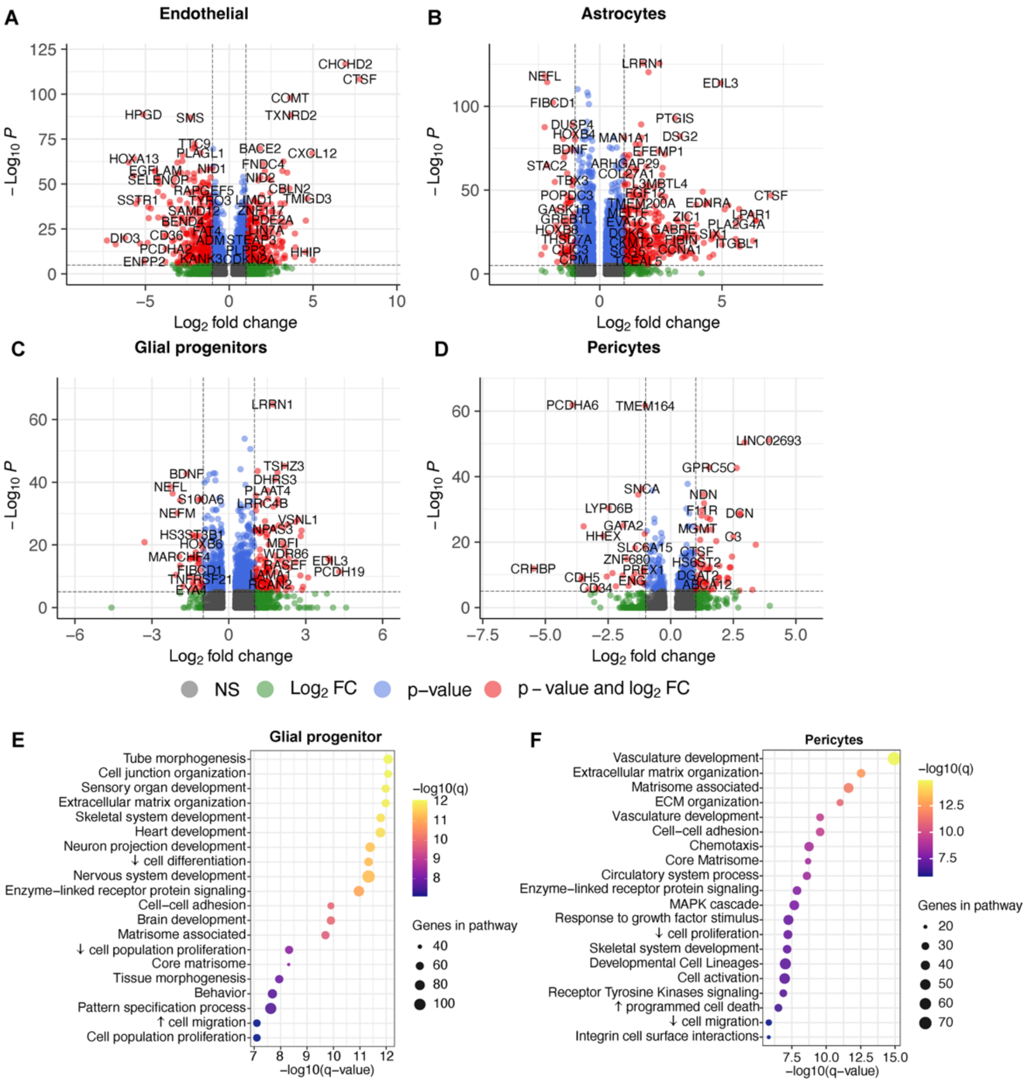


**Supplementary Figure 4. Differential expression for microtissue cell types.** Volcano plots summarising differential gene expression (MT vs corrected) for endothelial cells (**A**), astrocytes (**B**), glial progenitors (**C**), and pericytes (**D**). Genes are filtered and highlighted using |log2FC| ≥ 1 and P < 0.05 thresholds. Grey points indicate non-significant (ns) genes, green indicates genes meeting the |log2FC| cutoff only, blue indicates genes meeting the P-value cutoff only, and red indicates genes meeting both P-value and |log2FC| cutoffs. (**E,F**) Selected pathway-enrichment analyses in glial progenitor **(E)** and pericytes **(F)**.

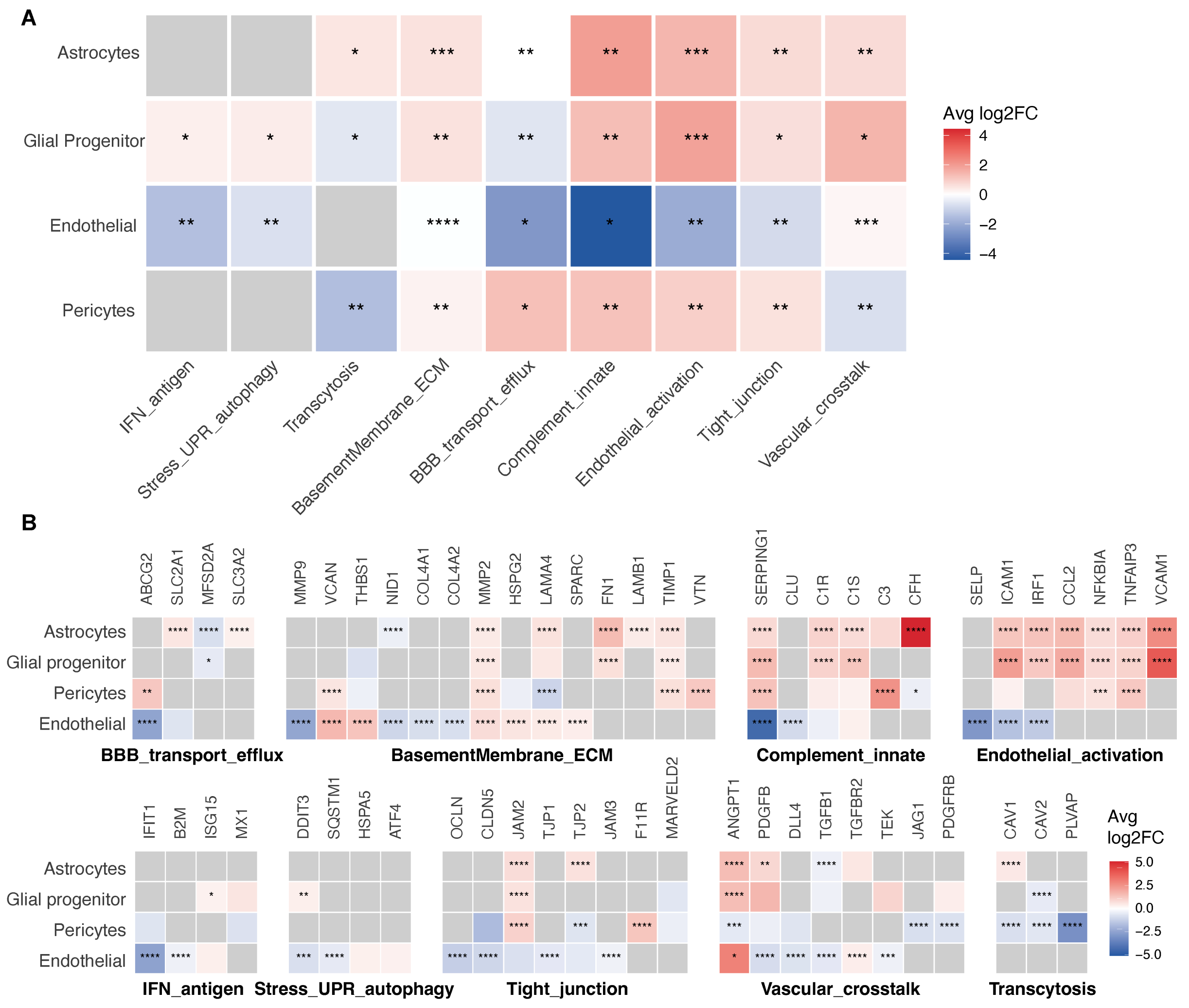


**Supplementary Figure 5. Cell-type-specific transcriptional changes in BBB, complement, extracellular matrix and inflammatory programmes.** (A) Heatmap showing mean log2 fold changes of curated functional gene modules across astrocytes, glial progenitor-like cells, endothelial cells and pericytes in MT relative to corrected BBB-like microtissues. Modules encompass BBB transport and efflux, tight junctions, transcytosis, complement-associated signalling, extracellular matrix (ECM) organisation, endothelial activation and related cellular processes. (B) Gene-level heatmaps showing differential expression of representative genes within each functional module across the four cell populations. Red and blue indicate increased and decreased expression in MT microtissues, respectively. Grey indicates undetected changes. *$p < 0.05$, **$p < 0.01$, ***$p < 0.001$, ****$p < 0.0001$.

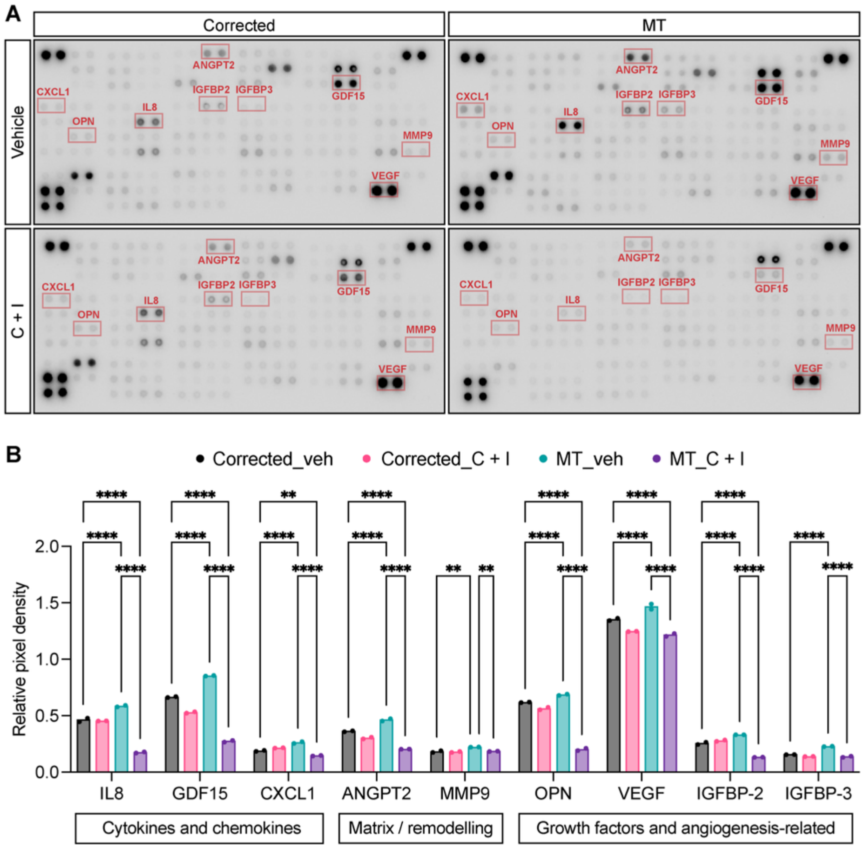


**Supplementary Figure 6. Exploratory secretome profiling following cilostazol and ISMN treatment. (A)** Representative human cytokine arrays of corrected and MT conditioned medium following vehicle or 1 µM cilostazol + 10 µM ISMN treatment. Conditioned media from 3 microtissues per group were pooled. **(B)** Relative spot intensities for selected inflammatory, matrix-related and angiogenesis-associated proteins. Data are presented as mean ± SEM, **p < 0.01, ***p < 0.001, ****p < 0.0001 by ordinary one-way ANOVA.

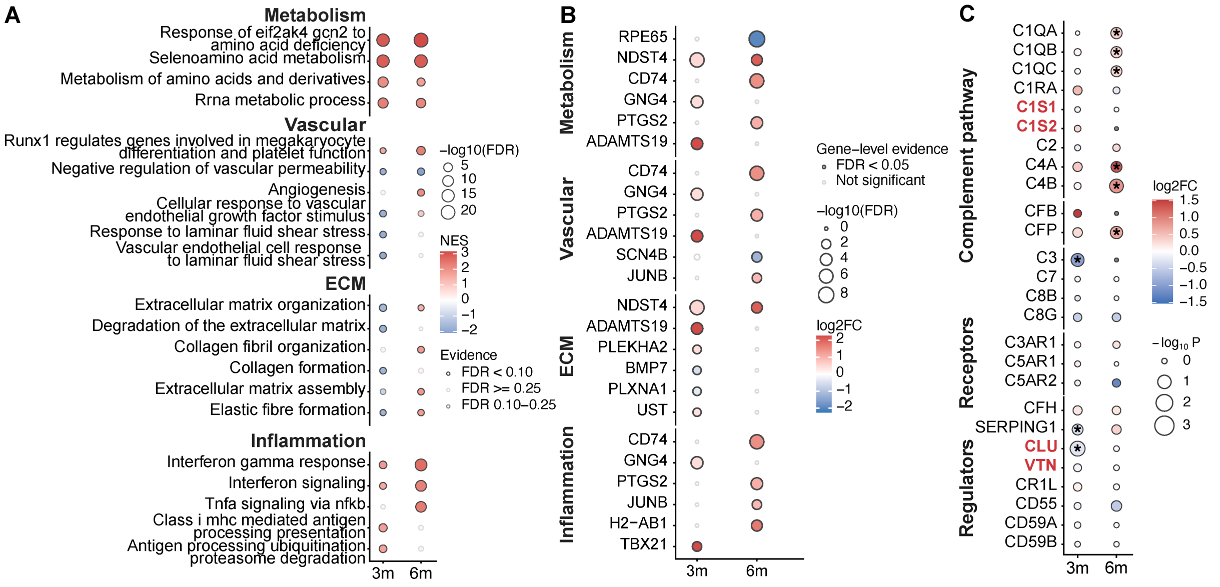


**Supplementary Figure 7. Age-associated transcriptomic and complement-related differences in NOTCH3 p.R544C mouse brain. (A)** Selected pathway enrichment across metabolism, vascular biology, extracellular matrix and inflammation. **(B,C)** Gene-level differences in representative metabolic/vascular and complement-associated programmes.

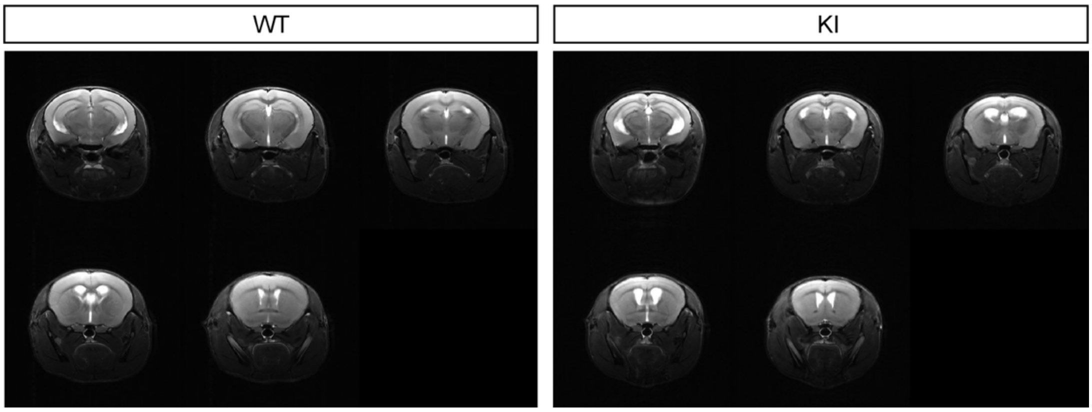


**Supplementary Figure 8. Preserved gross brain morphology in young NOTCH3 p.R544C mice.** Representative coronal structural magnetic resonance images of WT and NOTCH3$^{KI}$ mouse brains at three months of age, showing selected anatomical levels. No overt differences in gross brain morphology or ventricular structure were apparent between genotypes on visual inspection.

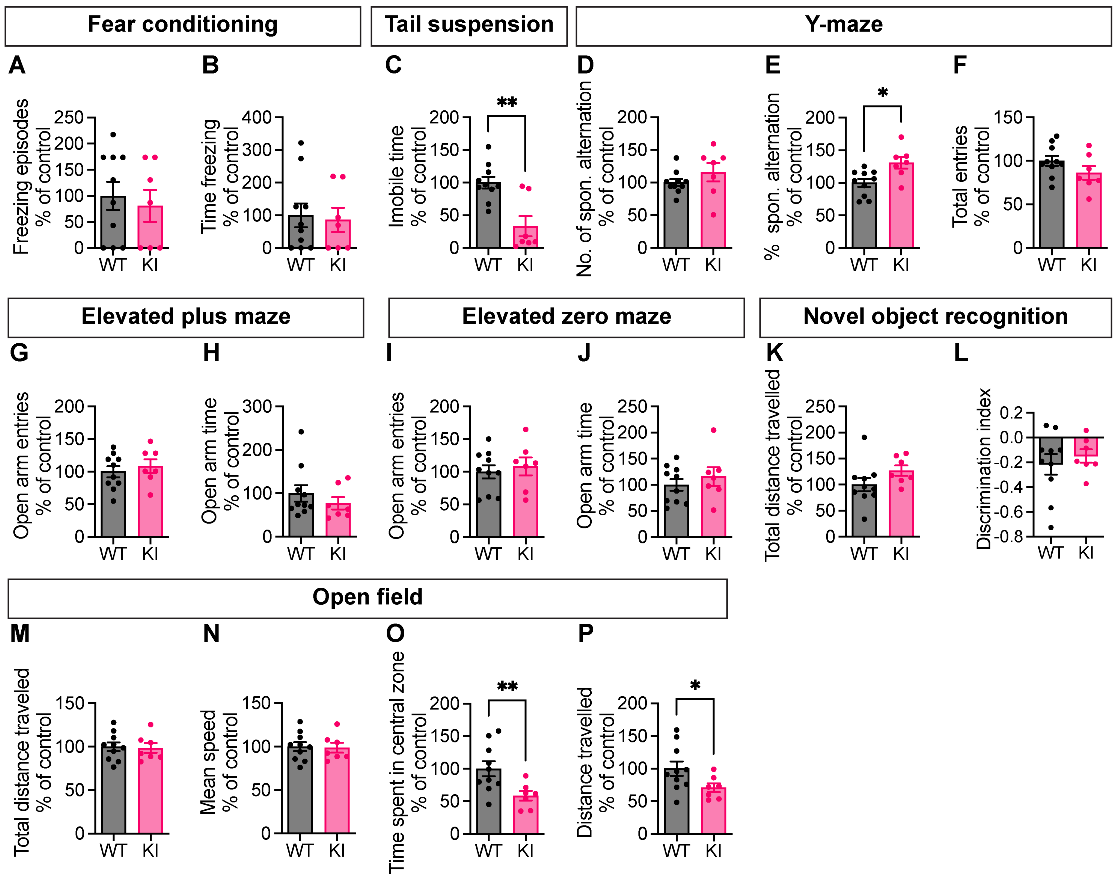


**Supplementary Figure 9. Behavioural measurements in three-month-old NOTCH3 p.R544C mice. (A,B)** Fear conditioning, **(C)** tail suspension, **(D-F)** Y-maze, **(G,H)** elevated plus maze, **(I,J)** elevated zero maze, **(K,L)** novel object recognition and **(M-P)** open-field testing in NOTCH3$^{KI}$ and WT mice. Data are presented as mean ± SEM, *p < 0.05, **p < 0.01 by Welch's t-test.
